# Microbiota-Based Interventions Differentially Rescue Gut and Social Behavior Phenotypes in a Drosophila Autism-like Model

**DOI:** 10.64898/2026.01.09.698713

**Authors:** Natalia A. Peta Martinez, Melanie Reinoso Arnaldi, Tasha M. Santiago-Rodriguez, Imilce A. Rodriguez-Fernandez

## Abstract

**Introduction:** Autism spectrum disorder (ASD) is a lifelong neurological and developmental disorder that has no cure and is often accompanied by gastrointestinal (GI) issues. The bidirectional communication system known as the gut microbiota-brain axis may help explain how GI dysfunction contributes to neurological symptoms. Loss-of-function mutations in the histone demethylases *KDM5A*, *KDM5B* or *KDM5C* are found in patients with intellectual disability and ASD. Previous studies using a *Drosophila* Kdm5 loss-of-function (*Kdm5^LOF^*) ASD-like model revealed gut microbial dysbiosis, reduced abundance of *Lactiplantibacillus plantarum*, and impaired social behavior. While *L. plantarum* supplementation rescued intestinal abnormalities, it did not restore social behavior.

**Methods:** Here, we evaluated multiple microbiota-based interventions, including probiotic supplementation with *Lactiplantibacillus plantarum*, *Lactobacillus helveticus*, their combination, and fecal microbiota transplantation (FMT), to determine their capacity to modulate gut microbial composition and behavior in *Kdm5^LOF^* flies. Gut bacterial abundance was quantified using colony-forming unit (CFU) assays and full-length 16S rRNA gene sequencing. Social behavior was assessed using the social distance assay, while anxiety-like behavior and locomotion were evaluated using the open field test. Gut-specific *Kdm5* knockdown was used to assess tissue-specific contributions to microbiota and behavioral phenotypes.

**Results:** *Kdm5* deficiency resulted in reduced abundance of culturable *Lactobacillus*, *Acetobacter*, and *Enterobacter* species, accompanied by impaired social behavior. *L. plantarum* supplementation restored gut microbial abundance in both whole-body *Kdm5^LOF^* and gut-specific *Kdm5* knockdown models but did not significantly rescue social behavior. In contrast, *L. helveticus* significantly improved social interaction in *Kdm5^LOF^* flies despite minimal effects on gut bacterial abundance, revealing a dissociation between microbial restoration and behavioral outcomes. Gut-specific *Kdm5* knockdown phenocopied both microbial and social defects observed in *Kdm5^LOF^*mutants. Notably, FMT from healthy donors partially restored *Lactobacillus* abundance, reshaped gut microbial community structure, and partially improved social behavior in *Kdm5^LOF^* recipient flies.

**Conclusions:** Together, these findings identify *Kdm5* as a key regulator of gut microbial viability and social behavior and demonstrate that microbiota-based interventions exert strain- and phenotype-specific effects. Our results reveal that restoration of microbial abundance alone is insufficient to rescue social behavior and highlight the importance of functional host–microbe interactions in gut–brain communication. This work establishes *Drosophila* as a tractable platform for dissecting epigenetic regulation of microbiota–behavior relationships relevant to ASD and for evaluating targeted probiotic and microbiota-transfer strategies.

## Introduction

Autism spectrum disorder (ASD) is a lifelong neurodevelopmental condition that emerges during early childhood [1]. The prevalence of ASD has increased substantially over the past decades, with recent estimates indicating that approximately 1 in 31 children have ASD in the United States [1]. ASD encompasses a wide spectrum of clinical presentations, including repetitive behaviors, restricted interests, sensory sensitivities, and impairments in social interaction and communication [2]. Twin and family studies have demonstrated a strong genetic contribution to ASD risk, with hundreds of genes implicated through rare and common variants that together contribute to a polygenic architecture [3]. Despite significant advances in genetic studies, the etiology of ASD remains incompletely understood, suggesting that multiple genetic and environmental factors interact during development to shape disease outcomes.

In addition to core behavioral features, ASD is frequently accompanied by comorbid medical conditions, most notably gastrointestinal (GI) dysfunction. Children with ASD are significantly more likely than neurotypical peers to experience GI symptoms such as constipation, diarrhea, abdominal pain, bloating, and vomiting [4,5]. Although these symptoms are highly prevalent, it remains unclear whether GI dysfunction contributes directly to neurobehavioral abnormalities or represents a parallel comorbidity [6]. Increasing attention has therefore focused on the gut microbiota–brain axis as a potential mechanistic link between intestinal physiology and neural development, with emerging evidence implicating microbial signals in neurodevelopmental processes relevant to ASD (reviewed in [7]).

The gut microbiota comprises a complex community of bacteria, fungi, archaea, and viruses that plays a critical role in host development, immune regulation, and metabolism. Communication between the gut and the central nervous system occurs through multiple pathways, including neural, immune, endocrine, and microbial metabolite–mediated signaling [7–9]. During development, this bidirectional communication is thought to influence neural circuit formation and behavioral outcomes [7]. Alterations in gut microbial composition have been associated with changes in neuroimmune signaling and inflammatory responses that are increasingly implicated in neurodevelopmental disorders, including ASD [7]. However, the molecular mechanisms and neural circuits linking microbial signals to behavior remain incompletely defined.

Numerous studies have reported differences in gut microbial diversity and composition in individuals with ASD compared to neurotypical controls, although specific microbial signatures vary across cohorts [5,7]. These observations have motivated the testing of microbiota-based interventions, including probiotic supplementation, aimed at improving GI and behavioral symptoms (reviewed in [8]. Probiotics are defined by the World Health Organization (WHO) as “live microorganisms that confer health benefits when administered in adequate amounts” [10]. In mouse ASD-like models, probiotic supplementation has been shown to improve certain behavioral phenotypes, including social deficits [11–14]. However, clinical and preclinical outcomes remain variable, and emerging evidence suggests that probiotic efficacy depends on factors such as host genotype, microbial community context, strain specificity, and developmental timing [8,15].

Beyond single-strain probiotics, fecal microbiota transplantation (FMT) has been explored as a community-level intervention to restore microbial diversity and function (reviewed in [16]). FMT is an established treatment for recurrent *Clostridioides difficile* infection and has shown promise in early clinical trials for ASD, with reported improvements in gastrointestinal symptoms and, in some cases, behavioral measures [17–20]. Nevertheless, the mechanisms underlying these effects remain poorly understood, and the extent to which microbiota replacement can overcome host genetic constraints on neurobehavioral phenotypes is unclear.

Simple animal model organisms provide a powerful framework for dissecting the interactions between host genetics, gut microbiota, and behavior [21]. The fruit fly *Drosophila melanogaster* has been extensively used to study neurodevelopment, behavior, and host–microbe interactions due to its genetic tractability and relatively simple gut microbiota (reviewed in [21–25]). Despite this simplicity, flies share conserved signaling pathways and microbial taxa with mammals, including bacteria from the Bacillota (formerly Firmicutes) and Pseudomonadota (formerly Proteobacteria) phyla, such as *Lactobacillus*, *Acetobacter*, and *Enterobacter* species [26,27]. Importantly, *Drosophila* enables precise manipulation of host genotype while controlling microbial exposure, allowing causal relationships between genes, microbiota, and behavior to be examined [22–24].

Mutations in chromatin regulators are among the most consistently identified genetic risk factors for ASD [28,29]. The KDM5 family of histone demethylases regulates transcription by removing methyl groups from histone H3 lysine 4 (H3K4), a modification associated with active gene expression [30]. In humans, loss-of-function mutations in KDM5A, KDM5B, and KDM5C are associated with ASD and intellectual disability [29,31–33]. *Drosophila* possesses a single ortholog, *Kdm5*, which has been shown to play essential roles in development, including neurodevelopment, synaptic structure, and neurotransmission [30,34,35].

Previous work by Chen et al. (2019) using a *Drosophila* ASD-like model carrying a loss-of-function mutation in *Kdm5* (hereafter referred to as *Kdm5^LOF^*) reported that disruption of this epigenetic regulator leads to significant alterations in gut microbial composition, reduced microbial diversity, and impaired intestinal barrier integrity [36]. These flies also exhibited pronounced social behavior defects, including increased social spacing and reduced contact time. Notably, supplementation with the commensal bacterium *Lactiplantibacillus plantarum* was sufficient to rescue intestinal defects but failed to fully restore social behavior [36]. These findings raise the possibility that while gut microbiota modulation can ameliorate physiological abnormalities, behavioral phenotypes may be influenced by host genetic and epigenetic context, and may not be fully responsive to single-strain probiotic supplementation.

We hypothesized that expanding microbiota-based interventions beyond single-strain probiotic supplementation, either through alternative probiotic species or broader microbial community replacement, would differentially affect intestinal and behavioral phenotypes in a genetically defined ASD-like model.

The present study builds on these findings by testing whether alternative microbiota-based strategies can overcome the limitations observed with single-strain supplementation. Specifically, we investigated whether supplementation with *Lactobacillus helveticus*, alone or in combination with *L. plantarum*, as well as fecal microbiota transplantation, could differentially modulate intestinal and behavioral phenotypes in *Kdm5* loss-of-function flies. *Lactobacillus helveticus* was selected based on prior evidence demonstrating its ability to modulate gut physiology, immune responses, and behavior across multiple animal models, including neurobehavioral contexts (reviewed in [8,37]). This allowed us to test whether probiotic effects on behavioral phenotypes in the *Kdm5* mutant model are strain-specific or reflect broader microbiota-driven mechanisms. By comparing targeted probiotic approaches with broader microbiota replacement, this work aims to define the extent to which microbiota manipulation can modulate gut physiology and behavior in a genetically defined ASD-like model, and to clarify the limits imposed by host epigenetic regulation on gut–brain interactions.

## Methods

### Fly husbandry and Fly stocks

Flies were raised on Nutri-Fly Food Bloomington-formula food (Genesee Cat. No. 66-113) and maintained at 25°C, >65% humidity and on a 12 h light/dark cycle, unless indicated otherwise. Each genetic scheme required multiple crosses to generate the experimental fly lines of interest.

From the Bloomington Drosophila Stock Center (BDSC), we obtained the following strains: *cn^1^, kdm5^10424^/CyO; ry^506^* (BDSC #12367) and *y^1^w^67c23^; kdm5^k06801^/CyO* (BDSC #10403). To remove unwanted genetic elements, these strains were crossed to additional balancer stocks, producing the intermediate lines *w^1118^; cn1, kdm5^10424^/CyO* and *w^1118^; kdm5^k06801^/CyO*. These two lines were subsequently crossed to generate a loss-of-function *Kdm5* (*Kdm5^LOF^*) mutant, (*w^1118^; kdm5^k6801^/kdm5^10424^,cn^1^*) as previously described and validated by Chen et al. (2019) [36]. For the genetic control, *w¹¹¹⁸* was crossed to *+; cn¹* (BDSC #263), because the *kdm5¹⁰⁴²⁴* chromosome carries a *cn¹* allele that could not be separated by recombination. All genotypes used for each experiment are summarized in **Supplementary Table 1**.

For the second genetic strategy, the primary fly stocks used were w*; *Myo31DF-Gal4; UAS-CC3Ai* (BDSC #84286); *y^1^v^1^*; *UAS-Kdm5^RNAi^* (P{TRiP.HM05155}attP2; BDSC #28944), *y^1^,v^1^; UAS-luciferase^RNAi^*(BDSC #31603) and *w*; esg-Gal4, UAS-2XYFP/CyO; Su(H)-GBEGal80, tub-Gal80^ts^/TM6* (gift from Heinrich Jasper, Genentech Inc). The *UAS-Kdm5^RNAi^*(P{TRiP.HM05155}attP2; BDSC #28944) was validated by [36]. Each stock was crossed to balancer lines as needed to isolate the required chromosomes and obtain the intermediate combinations. All experimental genotypes derived from these crosses are listed in **Supplementary Table 1**, in which *UAS-luciferase^RNAi^* served as the RNAi control. These parental crosses were maintained at 18°C to avoid RNAi expression during development. Adult flies of the desired genotype were collected, aged for 5–7 days, and subsequently shifted to 29°C for 3 days to induce gut-specific knockdown of *Kdm5 (*or induce *luciferase^RNAi^*, control) using the temperature-sensitive TARGET system (Gal4/Gal80^ts^) [38].

All experiments were conducted using mated female flies, as female midguts are larger, exhibit higher basal turnover rates, and display increased regenerative activity due to reproductive demands. Female flies are therefore the standard model for intestinal regeneration studies, following the discovery of midgut intestinal stem cells in 2006 [39,40].

### Probiotic Administration

From the parental cross, female progeny of the desired phenotypes were collected, allowed to mate for at least three days, and aged to 5–7 days. Flies were then divided into treatment groups based on the probiotic administered (**Table 1**). Probiotics were applied directly onto the surface of the food, and flies fed *ad libitum* for 2–3 days before downstream assays.

**Table 1.**
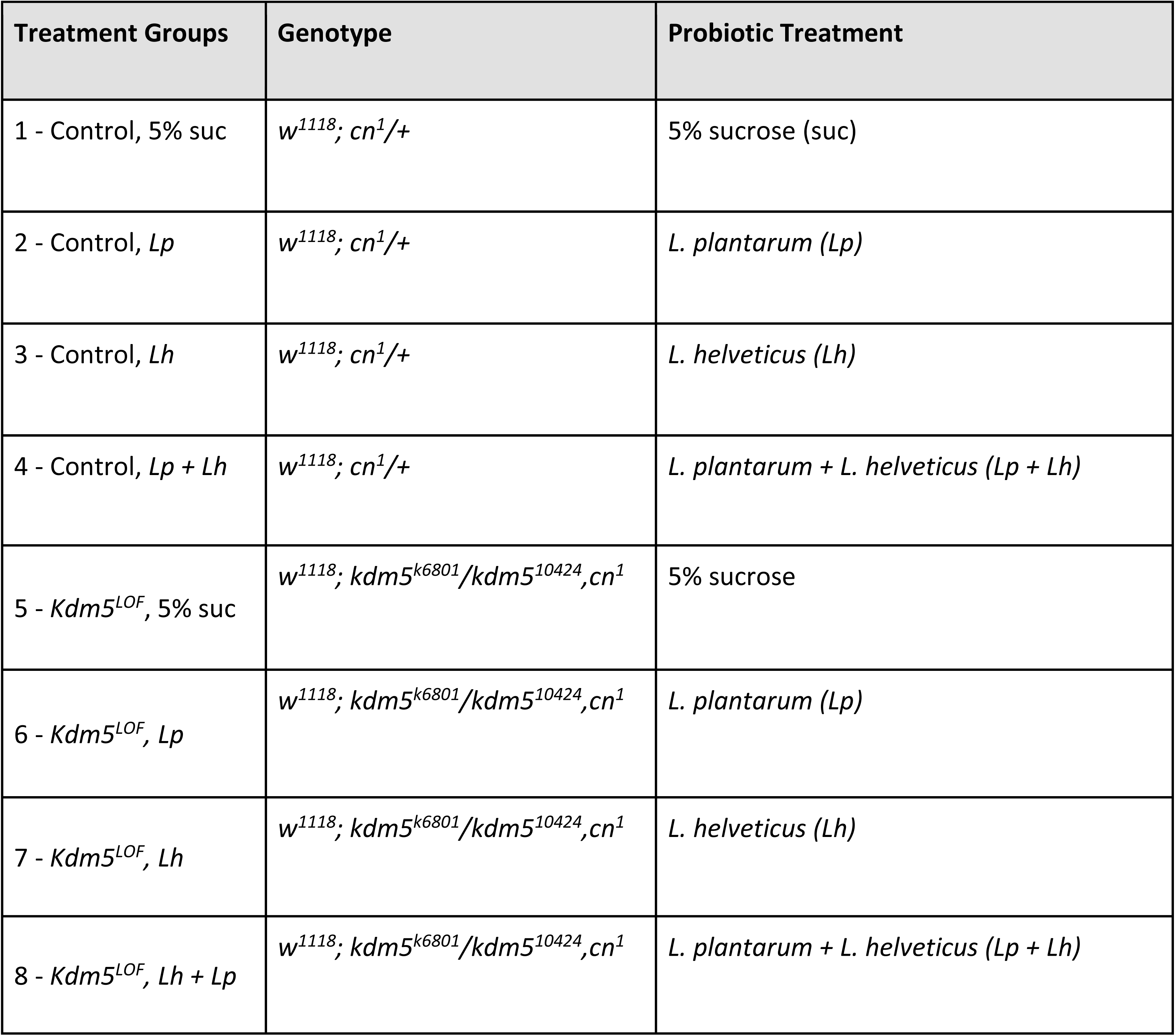
Assignment of some treatment groups used by genotype and probiotic treatment.

The probiotics used were *Lactiplantibacillus plantarum* (formerly known as *Lactobacillus plantarum*) (ATCC 14917) and *Lactobacillus helveticus* (ATCC 15009), both obtained from the American Type Culture Collection (ATCC). From 20% glycerol stocks, bacteria were inoculated on MRS broth (BD Difco™ Lactobacilli MRS Broth BD288130) and incubated at 37°C, shaking at 200 rpm, for 24–48 hours. Cultures were then centrifuged at 4,000 rpm for 10 minutes at 25°C, and the resulting pellets were resuspended in sterile 5% sucrose (w/v) prepared in distilled water. Mock control contains only 5% sucrose.

Bacterial density was measured at OD₆₀₀ using a spectrophotometer (Bio-Rad). For all experiments, the final working concentration of bacteria was standardized to 1 × 10⁸ cells/mL. A 200 µL aliquot of this bacterial suspension was applied onto the surface of food vials (containing ∼ 10 mL of food) that had been pre-punctured with a sterile syringe to facilitate absorption. Vials were prepared approximately 1 hour before introducing the flies to allow sufficient time for the suspension to soak into the food. Flies were maintained on these probiotic-supplemented vials for 2–3 days, as indicated, prior to phenotypic assessments.

### Colony Forming Unit

The colony-forming unit (CFU) assay was used to quantify viable bacterial cells in dissected fly guts and was performed as previously reported [41,42]. Three types of selective agar plates were used: *Lactobacilli* MRS agar, *Acetobacter* agar, and *Enterobacter* agar. BD Difco™ MRS agar was obtained from Fisher Scientific (Cat. No.: BD 288130). *Acetobacter* agar was prepared by dissolving 25 g/L D-mannitol (CAS No.: 69-65-8; Avantor Sciences VWR Cat. No.: BDH9248), 5 g/L yeast extract (Gibco™ Bacto™ Yeast Extract; Fisher Scientific Cat. No.: Gibco 212750), 3 g/L peptone (Fisher Scientific Cat. No.: OXLP0037B), and 15 g/L agar (Fisher Scientific Cat. No.: BP1423-500) in distilled water.

*Enterobacter* agar was prepared using 10 g/L tryptone (Gibco™ Bacto™ Tryptone; Fisher Scientific Cat. No.: Gibco 211705), 1.5 g/L yeast extract, 10 g/L glucose (CAS No.: 50-99-7; Millipore Sigma Cat. No.: G7021), 5 g/L NaCl, and 12 g/L agar. All media were autoclaved at 121°C for 15 min, poured into plates, and stored at 4°C. Before use, plates were equilibrated to room temperature and then warmed briefly at 37°C.

All dissections and sample preparation steps were performed under sterile conditions. Work surfaces were cleaned with 70% ethanol, allowed to dry, and maintained sterile using a Bunsen burner flame. Flies were surface-sterilized by sequential immersion in 70% ethanol for 30 seconds, followed by two rinses in sterile 1x Phosphate Buffered Saline (PBS) of 10 seconds each. Dissections were performed in a drop of sterile PBS on sterile Petri dishes (60 mm x 15 mm) (Fisher Scientific Cat. No.: AS4052).

Dissected guts were transferred to sterile microcentrifuge tubes containing 200 µL of sterile PBS and homogenized using a sterile pestle (Fisher Scientific Cat. No.: 12141368) attached to a hand-held homogenizer (Avantor Biosciences VWR® Cordless Pestle Motor, Cat. No. 47747-370) until no visible tissue remained. An additional 300 µL of sterile PBS was added to bring the final volume to 500 µL. Homogenates were serially diluted in sterile PBS to 1:100 and 1:1000.

For plating, 2 µL of each dilution was spotted onto the appropriate agar plates in separate, non-overlapping spots, with three drops plated per condition as technical replicates. Plates were incubated at 37°C and examined after 24–48 hours. Visible bacterial colonies (microcolonies) were counted to determine CFUs per gut. As a negative control, sterile PBS used during the experiment was also spotted onto plates to confirm the absence of contamination. In all experiments, no bacterial growth was detected from PBS controls.

### Social Distance Assay

The social distance assay measures social space, defined as the distance between an individual fly and its nearest neighbor, and was adapted from the protocol described by [43]. The assay was performed using a custom-built vertical chamber composed of two square glass panes (17.6 × 17.6 cm, 0.3 cm thickness), two right-triangle acrylic spacers (16.5 cm height, 8.9 cm base, 0.3 cm thickness), and two rectangular acrylic spacers (9 × 1.5 cm, 0.3 cm thickness). To assemble the chamber, one glass pane was placed flat on the bench, and the two right-triangle spacers were positioned along the left and right edges, meeting at the top center to form an internal triangular cavity with a bottom distance of 15.5 cm. The rectangular spacers were placed along the bottom edge of the glass pane. A second glass pane was then aligned on top, and the chamber was secured using four 5-cm binder clips (one per corner).

Between 13-55 young (5-7 days) female flies of the indicated genotype and treatment were collected for the experiment. On the day of the experiment, these flies were placed in the room where the assays were done for 1 hour before starting the social distance experiment. When this time passed, flies were briefly cold-anesthetized by transferring them to empty vials and placing the vials at a −20°C freezer for approximately 2 minutes 15 seconds (maximum). Following cold anesthesia, flies were introduced into the chamber by removing one bottom binder clip and gently displacing a rectangular spacer to create a small opening. Flies were transferred using a homemade mouth aspirator and gently expelled into the chamber to promote aggregation into a tight group. The spacer was immediately returned to its original position and secured with the binder clip. One treatment/condition was done at a time.

To standardize the starting position, the chamber was placed vertically and tapped firmly against a padded surface (mouse pad) to allow flies to aggregate at the bottom of the chamber. The chamber was then maintained in a vertical position, and flies were allowed to acclimate for 10 minutes. After acclimation, the vertically oriented chamber was photographed against a white background using an iPhone smartphone (model 11) 10 minutes after acclimatization.

Images were analyzed in ImageJ/FIJI [44]. The scale of the triangular chamber was first set, after which individual flies were manually marked using the multi-point tool and defined as regions of interest (ROIs). Euclidean distances between each fly and its nearest neighbor were calculated from the ROI coordinates. Distances (in cm) were exported and plotted as histograms in Microsoft Excel using a bin width of 0.5 cm. The Social Space Index (SSI) was calculated as described by [43], defined as the percentage of flies within the first distance bin minus the percentage within the second bin (SSI = %Bin₁ – %Bin₂). SSI values ≤ 0 indicate reduced social interaction.

Statistical analyses of SSI were performed using GraphPad Prism. A two-way ANOVA followed by Šídák’s or Dunnetts multiple-comparisons test (as indicated in figure legend) was used to assess the effects of genotype and treatment on social distance. Each experiment was performed independently three times (three biological replicates), with each replicate consisting of one technical replicate of 13–55 flies.

### Fecal Microbiota Transplantation (FMT)

Fecal microbiota transplantation (FMT) was performed to assess the impact of donor-derived gut microbiota on intestinal bacterial composition and behavior in recipient flies. Adult control and *Kdm5^LOF^*flies (5–7 days old) were used as both fecal donors and recipients.

For donor conditioning, flies were placed in standard food vials and maintained for 3 days to allow accumulation of fecal material. For experiments followed by the social distance assay, approximately 40 donor flies were placed per vial, whereas for CFU-based analyses, 15–20 donor flies were used per vial. Donor flies were then removed, leaving behind food conditioned with fecal material.

Recipient flies (5–7 days old) were subsequently transferred into the conditioned vials and exposed to donor fecal material for 24 h. Four donor–recipient combinations were generated: control recipients exposed to control donor feces, *Kdm5^LOF^*recipients exposed to *Kdm5^LOF^*donor feces, control recipients exposed to *Kdm5^LOF^* donor feces, and *Kdm5^LOF^* recipients exposed to control donor feces. During this period, recipient flies ingested donor-associated microbiota through feeding on the conditioned food.

Following the 24-h exposure period, recipient flies were collected and processed for downstream analyses, including CFU quantification and behavioral assays, as indicated.

### 16S rRNA gene sequencing and microbiome analysis following FMT

Following FMT experiments, ten flies per genotype were dissected under axenic conditions as described in the ‘Colony Forming Units’ section. For each biological replicate, ten female flies per genotype (5–7 days old) were dissected, and guts were pooled in sterile microcentrifuge tubes on ice containing cold PBS. After collection, PBS was removed by pipetting, and samples were flash-frozen on dry ice and stored at −80 °C until processing. Each experimental condition consisted of three independent biological replicates, each comprising ten pooled guts.

Frozen samples were shipped on dry ice and processed by the ZymoBIOMICS® Targeted Sequencing Service (Zymo Research, Irvine, CA). Genomic DNA was extracted using either the ZymoBIOMICS®-96 MagBead DNA Kit or the ZymoBIOMICS® DNA Miniprep Kit (Zymo Research, Irvine, CA). The MagBead kit was used for most samples and processed on an automated platform, whereas the Miniprep kit was used for low-biomass samples to allow lower elution volumes and increased DNA concentration.

Full-length 16S rRNA gene libraries were prepared following the PacBio full-length 16S amplification protocol. Briefly, the 16S rRNA gene was amplified using barcoded universal primers 27F (AGRGTTYGATYMTGGCTCAG) and 1492R (RGYTACCTTGTTACGACTT). For each sample, 2 ng of DNA was used as a PCR template, and amplification was performed for 25 cycles under conditions specified in the protocol. Amplicons were purified using the Select-a-Size DNA Clean & Concentrator MagBead Kit (Zymo Research, Irvine, CA), retaining fragments >300 bp. Purified libraries were quantified using NanoDrop, pooled at equimolar concentrations, and converted into SMRTbell® libraries using the SMRTbell® Prep Kit 3.0 (PacBio).

Positive controls included the ZymoBIOMICS® Microbial Community Standard for DNA extraction and the ZymoBIOMICS® Microbial Community DNA Standard for library preparation. Negative controls, including blank extraction and blank library preparation controls, were included to assess potential background contamination. Sequencing was performed on a single 8M SMRT Cell using the PacBio Sequel IIe system.

Raw sequencing reads were processed using the DADA2 pipeline to infer amplicon sequence variants (ASVs), remove sequencing errors, and filter chimeric sequences. Taxonomic assignment was performed using UCLUST within QIIME v1.9.1, using the Zymo Research curated 16S reference database. Alpha diversity was evaluated using observed taxon richness, the Shannon diversity index, and the Inverse Simpson index, calculated in RStudio (v. 2025.09.02+148) with the phyloseq package (v. 1.54.0). Samples were rarefied to an even sequencing depth using the rarefy_even_depth function without replacement. Statistical significance (p < 0.05) was assessed using the Kruskal-Wallis test. Alpha diversity metrics were visualized using boxplots generated with ggplot2 (v. 4.0.1). Beta diversity was assessed using the Agile Toolkit for Incisive Microbial Analyses (ATIMA; https://atima.research.bcm.edu/) based on Bray–Curtis dissimilarity metrics. The uploaded taxonomy table was rarefied to 23,544 reads per sample, and community differences were visualized using Principal Coordinates Analysis (PCoA) plots. Statistical significance of group differences was evaluated using permutational multivariate analysis of variance (PERMANOVA). Relative abundances of the identified taxa were visualized through stacked bar plots generated using the package MicrobeR (v.0.3.2) and the function Microbiome.Barplot. Differential abundance analyses were performed using the package MaAsLin2 in RStudio with Total Scale Sum (TSS) normalization and log transformation. Results were then filtered to visualize differentially abundant taxa with q-value < 0.05 using lollipop plots generated using ggplot2.

## Results

### *L. plantarum* rescues microbial dysbiosis in *Kdm5^LOF^* flies, whereas *L. helveticus* shows no effect

To assess microbial dysbiosis in female *Drosophila* treated as described in **Table 1**, we performed CFU assays. CFU analysis is a well-established method to quantify viable gut-associated bacteria and is commonly used to evaluate alterations in microbial community structure in *Drosophila* models of intestinal dysfunction and disease [41,42,45–47].

The CFU assay was used to quantify the abundance of the three major bacterial genera typically associated with the *Drosophila* gut microbiota: *Lactobacillus*, *Acetobacter*, and *Enterobacter*. Following treatment, adult female fly guts were dissected under sterile conditions, homogenized in sterile PBS, and plated as serial dilutions on selective media. After 24–48 hours of incubation, bacterial colonies were counted, and CFUs were calculated per gut.

CFU analysis revealed genus-specific alterations in gut microbial composition in *Kdm5^LOF^*flies compared to controls (**Figure 1A–C**). In control flies, the abundance of *Acetobacter* was not significantly affected by supplementation with *L. plantarum* (Lp), *L. helveticus* (Lh), or the combined treatment (Lp+Lh) (**Figure 1A**). In contrast, *Kdm5^LOF^* flies fed a 5% sucrose control diet exhibited a significantly reduced abundance of *Acetobacter* relative to control flies fed 5% sucrose. Supplementation with Lh did not alter *Acetobacter* levels in *Kdm5^LOF^* flies, whereas treatment with Lp or Lp+Lh significantly increased *Acetobacter* abundance, restoring levels toward those observed in control flies.

**Fig. 1.**
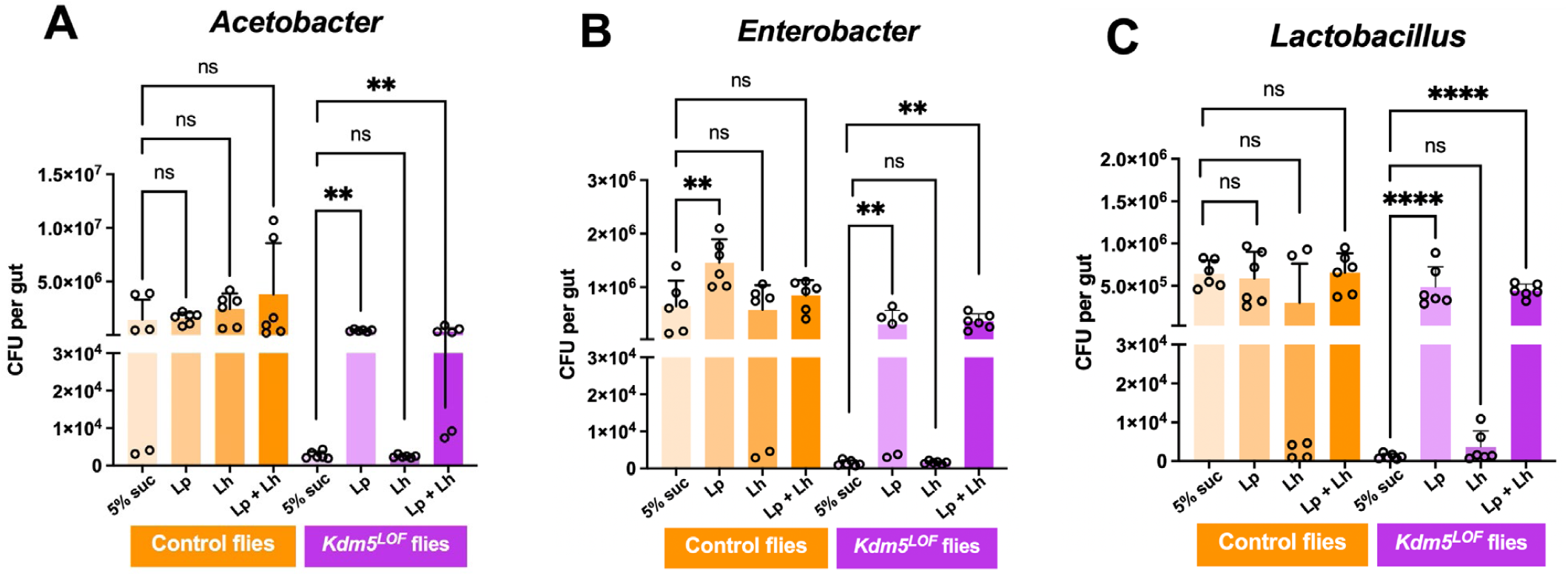
*L. plantarum* rescues microbial dysbiosis in *Kdm5^LOF^* flies. Young adult control and *Kdm5^LOF^* flies (5–7 days old) were treated with 5% sucrose (mock control), *Lactiplantibacillus plantarum* (Lp), *Lactobacillus helveticus* (Lh), or a 1:1 combination of Lp and Lh in 5% sucrose for 2–3 days, supplemented on standard fly food. Guts were dissected, homogenized and plated on selective agar media to quantify CFUs per gut of *Acetobacter* (A)*, Enterobacter* (B) and *Lactobacilli* (C) genera. Bars represent mean ± SEM from three independent biological replicates, with duplicate technical platings per sample. Statistical significance was determined by one-way ANOVA followed by Dunnetts multiple-comparisons test. **p < 0.01; ****p < 0.0001; ns, not significant.

A similar pattern was observed for *Enterobacter* (**Figure 1B**). In control flies, *Enterobacter* abundance was largely unchanged following Lh or Lp+Lh supplementation, although Lp treatment resulted in a significant increase. *Kdm5^LOF^* flies displayed a marked reduction in *Enterobacter* abundance compared to controls, which was not rescued by Lh treatment. In contrast, supplementation with Lp or Lp+Lh significantly increased *Enterobacter* abundance in *Kdm5^LOF^* flies.

Analysis of *Lactobacillus* species cultured on MRS agar revealed comparable trends (**Figure 1C**). Control flies maintained stable *Lactobacillus* levels across all treatment conditions. In *Kdm5^LOF^* flies, *Lactobacillus* abundance was significantly reduced relative to controls and was not restored by Lh supplementation. However, treatment with Lp or Lp+Lh resulted in a significant increase in *Lactobacillus* abundance.

Survival and intestinal barrier integrity were assessed using the Smurf assay for 14 days following mock or probiotic treatments. Survival remained high across all genotypes and treatment groups, and no Smurf phenotype was observed during the monitoring period (0/37 flies per group; **Supplementary Fig. 1**). Consistent with this, no significant differences were detected in either survival or gut permeability, indicating that the microbial dysbiosis observed in *Kdm5^LOF^* flies occurs independently of detectable changes in intestinal barrier integrity under the conditions tested.

### *L. helveticus* rescues social behavior deficits in *Kdm5^LOF^* flies, whereas *L. plantarum* shows no effect

To assess social behavior, we performed the social distance assay, which quantifies social space, defined as the distance between an individual fly and its nearest neighbor. In this assay, socially interacting flies cluster closely, whereas reduced social interaction is reflected by increased spacing between individuals. Distances to the nearest neighbor were measured and plotted as histograms representing the percentage of flies occupying 0.5-cm distance bins (**Figure 2A–H**). Social behavior was further quantified using the Social Space Index (SSI), calculated as the percentage of flies in the first distance bin minus the percentage in the second bin (**Figure 2I**). SSI values greater than zero indicate increased social proximity, whereas values at or below zero indicate reduced social interaction.

**Fig. 2.**
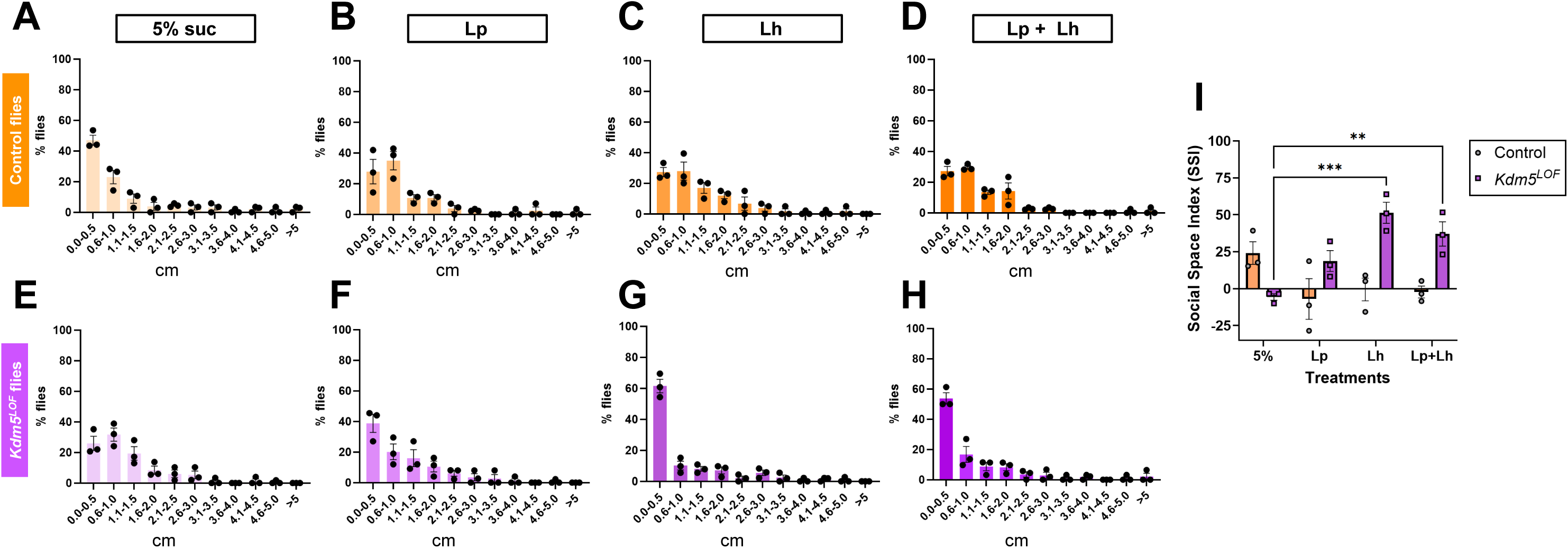
Probiotic treatment modulates social spacing behavior in *Kdm5^LOF^* and control flies. Young adult control and *Kdm5^LOF^* flies (5–7 days old) were treated with 5% sucrose (mock control), *Lactiplantibacillus plantarum* (Lp), *Lactobacillus helveticus* (Lh), or a 1:1 combination of Lp and Lh in 5% sucrose for 2–3 days, prior to behavioral testing. Social spacing behavior was assessed using the Social Distance assay. Histograms (A–H) show the percentage of flies occupying 0.5-cm distance bins, calculated based on the distance from the nearest to the farthest neighbor. Panels A–D correspond to control flies, and panels E–H correspond to *Kdm5^LOF^* flies under the indicated treatment conditions. (I) The Social Space Index (SSI) was calculated as the percentage of flies in the first distance bin minus the percentage in the second bin for each condition. Positive SSI values indicate increased social proximity, whereas values at or below zero indicate reduced social interaction. Bars represent mean ± SEM from three independent biological replicates. Statistical significance was determined using a two-way ANOVA followed by Šídák’s multiple-comparisons test. All post hoc comparisons were performed within genotypes; only the two statistically significant comparisons are indicated (**p < 0.01; ***p < 0.001).

Control flies exhibited robust social behavior under mock conditions (5% sucrose), characterized by a high proportion of flies occupying the first distance bin and minimal representation in subsequent bins (**Figure 2A**). In contrast, administration of probiotic treatments to control flies resulted in a redistribution of flies across larger distance bins (**Figure 2B–D**), suggesting a modest reduction in social proximity (**Figure 2I)**, although differences in SSI values were not significantly different from control 5% mock treated flies when analyzed using a two-way ANOVA followed by Šídák multiple-comparisons test.

*Kdm5^LOF^* flies displayed impaired social behavior under mock conditions, with a reduced proportion of flies in the first distance bin and an increased proportion in the second bin, consistent with diminished social interaction (**Figure 2E**). Treatment of *Kdm5^LOF^* flies with *L. plantarum* (Lp), *L. helveticus* (Lh), or a combination of both probiotics shifted the distribution toward increased occupancy of the first distance bin (**Figure 2F–H**), indicating a tendency toward increased social proximity. Statistical analysis of SSI values using a two-way ANOVA revealed a significant genotype × treatment interaction, indicating that probiotic effects on social behavior differed between genotypes. Post hoc Šídák multiple-comparisons tests were performed within each genotype. In *Kdm5^LOF^* flies, treatment with Lh and with the Lp + Lh combination resulted in a significant increase in SSI compared to mock-treated *Kdm5^LOF^* flies (**Figure 2I**), consistent with a rescue of the social behavior defect. No other within-genotype comparisons reached statistical significance.

Importantly, these effects were specific to social behavior, as open field testing revealed no significant differences in zone occupancy across genotypes or probiotic treatments, indicating that general locomotor activity and exploratory behavior were unaffected (**Supplementary Fig. 2**).

### Gut-specific knockdown of *Kdm5* phenocopies whole-body *Kdm5^LOF^*, with intestinal microbial dysbiosis rescued by *L. plantarum* but not *L. helveticus*

To determine whether intestinal reduction of Kdm5 is sufficient to recapitulate microbial phenotypes observed in whole-body Kdm5 loss-of-function flies, we quantified gut bacterial abundance in flies with adult, gut-specific Kdm5 knockdown (*Kdm5^RNAi^* using MyoD31-Gal4 driver) and corresponding *luciferase^RNAi^* controls following probiotic treatment. CFUs were quantified for *Acetobacter*, *Enterobacter*, and *Lactobacillus* genera (**Figure 3**).

**Fig. 3.**
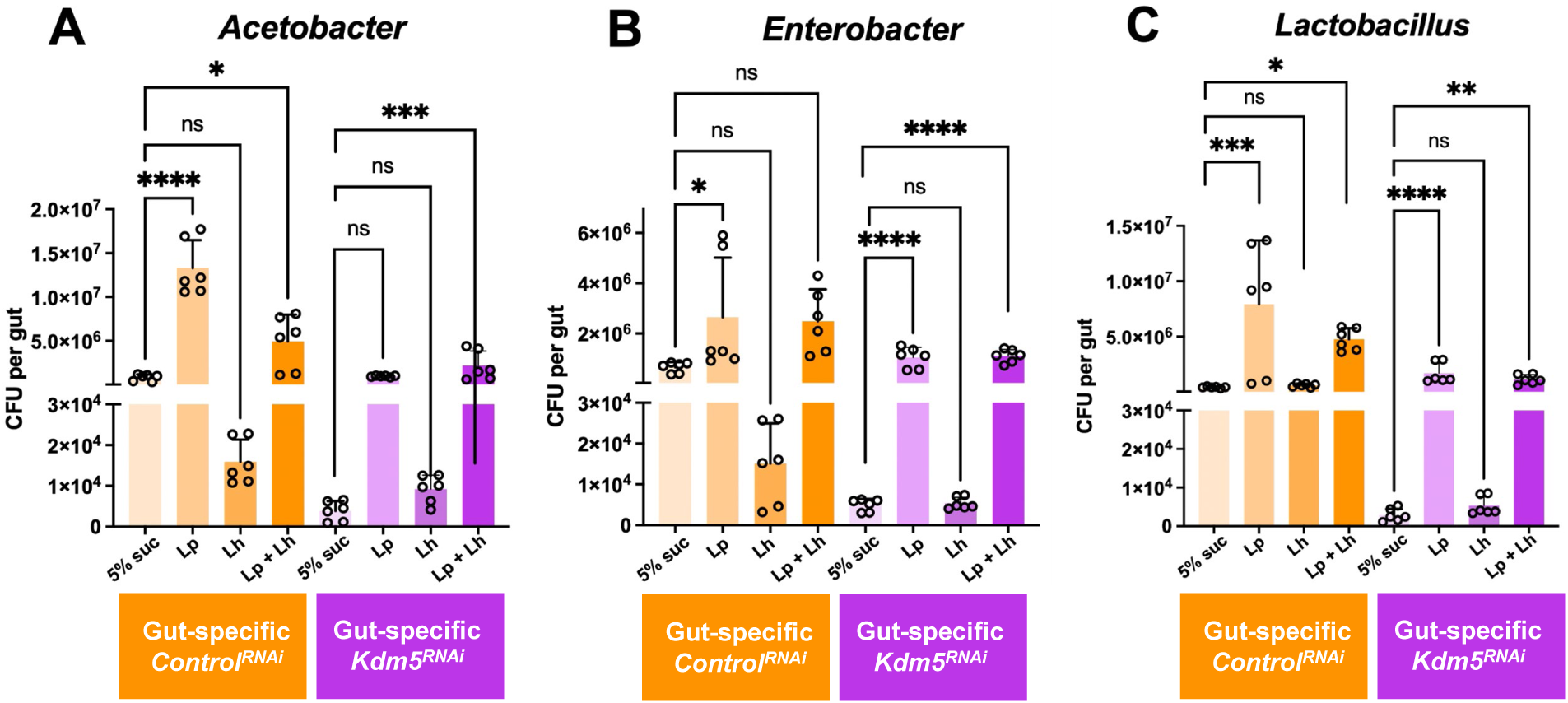
Gut-specific knockdown of Kdm5 alters intestinal bacterial abundance in a strain-dependent manner. Young adult control (gut-specific *Lucierase^RNAi^*) and gut-specific *Kdm5^RNAi^* flies (5-7 days old; RNAi induced at 29°C for 3 days) were treated with 5% sucrose (mock control), *L. plantarum* (*Lp*), *L. helveticus* (*Lh*) or a 1:1 combination of *Lp* and *Lh* in 5% sucrose for 2-3 days, supplemented on standard fly food. Guts were dissected, homogenized and plated on selective agar media to quantify CFUs per gut of *Acetobacter* (A)*, Enterobacter* (B) and *Lactobacilli* (C) genera. Bars represent mean ± SEM from three independent biological replicates, with duplicate technical platings per sample. Statistical significance was determined by one-way ANOVA followed by Dunnett’s or Šídák’s multiple-comparisons test, as indicated. *p < 0.05; **p < 0.01; ***p < 0.001; ****p < 0.0001; ns, not significant.

Control flies expressing luciferase RNAi exhibited stable and balanced gut microbial profiles across all three bacterial genera under mock conditions (5% sucrose) (**Figure 3A–C**). In these control flies, treatment with Lp or the Lp + Lh combination significantly increased *Acetobacter* abundance, whereas Lh alone did not (**Figure 3A**). For *Enterobacter*, Lp treatment resulted in a significant increase in abundance, while Lh showed a tendency toward reduced levels (**Figure 3B**). In contrast, *Lactobacillus* abundance in control flies was significantly increased following Lp or Lp + Lh treatment, with no significant change observed following Lh treatment alone (**Figure 3C**).

Gut-specific *Kdm5^RNAi^* flies exhibited reduced abundance of *Acetobacter*, *Enterobacter*, and *Lactobacillus* under mock conditions compared to controls, indicating intestinal dysbiosis from reducing Kdm5 levels specifically in the gut (**Figure 3A–C**). Probiotic treatment partially rescued these defects in a strain-dependent manner. The Lp + Lh combination significantly increased *Acetobacter* levels, whereas neither Lp nor Lh alone produced a significant effect (**Figure 3A**). For *Enterobacter*, both Lp and the Lp + Lh combination significantly increased bacterial abundance, while Lh alone failed to restore levels (**Figure 3B**). Similarly, *Lactobacillus* abundance was significantly increased by Lp and by the Lp + Lh combination, but not by Lh treatment alone (**Figure 3C**).

### Gut-specific Kdm5 knockdown leads to impaired social behavior that is selectively rescued by probiotic treatment

To assess whether gut-specific loss of *Kdm5* also impacts social behavior, we performed the social distance assay on *Kdm5^RNAi^* flies and *luciferase^RNAi^*controls following probiotic treatment (**Figure 4**). Social behavior was quantified using histograms of nearest-neighbor distances and summarized using the SSI.

**Fig. 4.**
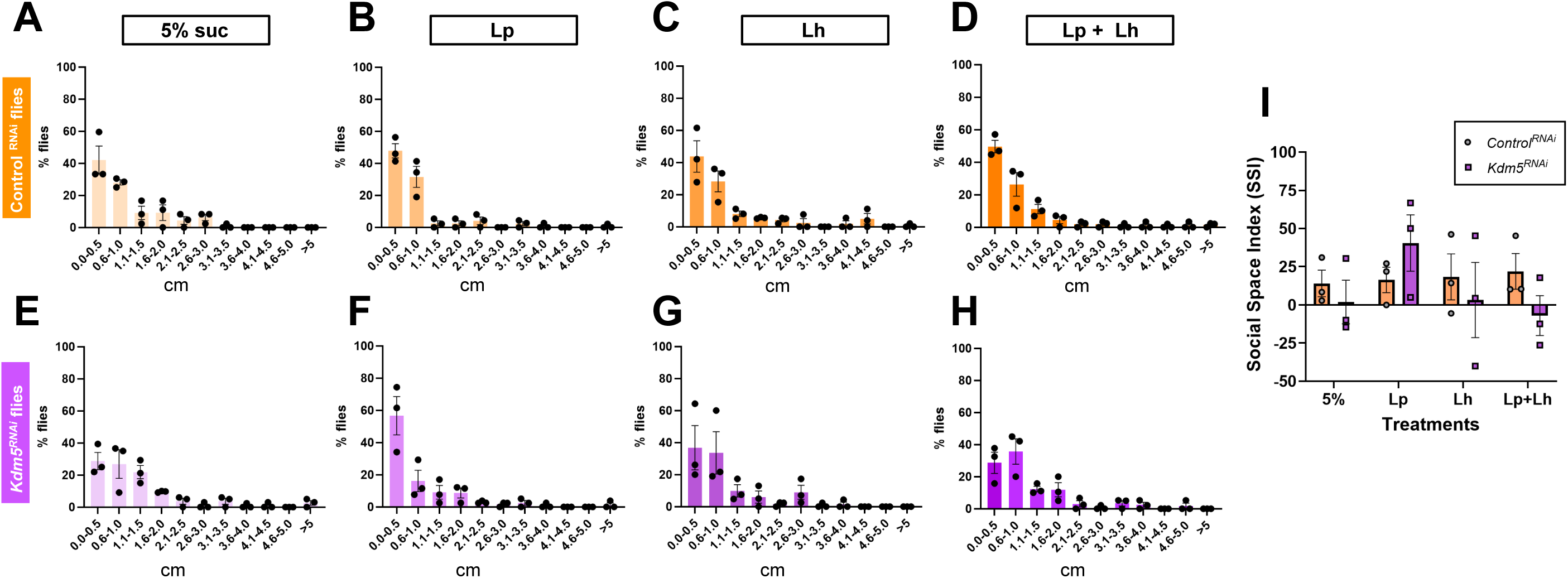
Gut-specific knockdown of Kdm5 alters social spacing behavior. Young adult control (gut-specific *Lucierase^RNAi^*) and gut-specific *Kdm5^RNAi^*flies (5-7 days old; (5-7 days old; RNAi induced at 29°C for 3 days) were treated with 5% sucrose (mock control), *L. plantarum* (*Lp*), *L. helveticus* (*Lh*) or a 1:1 combination of *Lp* and *Lh* in 5% sucrose for 2-3 days prior to behavioral testing. Social spacing behavior was assessed using the social distance assay. Histograms (A–H) show the percentage of flies occupying 0.5-cm distance bins, calculated based on the distance from each fly to its nearest neighbor. Panels A–D correspond to control flies, and panels E–H correspond to gut-specific *Kdm5^RNAi^* flies under the indicated treatment conditions. (I) The Social Space Index (SSI) was calculated as the percentage of flies in the first distance bin minus the percentage in the second bin for each condition. Positive SSI values indicate increased social proximity, whereas SSI values ≤ 0 indicate reduced social interaction. Bars represent mean ± SEM from three independent biological replicates. Statistical significance was determined using a two-way ANOVA followed by Šídák’s multiple-comparisons test; no post hoc SSI comparisons reached statistical significance.

*luciferase^RNAi^* control flies displayed robust social behavior under mock conditions, characterized by a high proportion of flies occupying the first distance bin (**Figure 4A**). Unlike the control flies used in the whole-body *Kdm5^LOF^*model (**Figure 2A-D**), probiotic treatments did not negatively impact social behavior in *luciferase^RNAi^*controls, and SSI values remained positive across all treatments (**Figure 4I**).

In contrast, gut-specific *Kdm5^RNAi^* flies exhibited impaired social behavior under mock conditions, with a shift toward increased occupancy of the second distance bin and reduced SSI values (**Figure 4E,I**), indicating diminished social interaction. Administration of probiotics resulted in a redistribution toward shorter inter-fly distances (**Figure 4F–H**). Although Lp produced a qualitative shift toward increased social proximity in gut-specific *Kdm5^RNAi^* flies compared to mock-treated animals, these effects did not reach statistical significance by two-way ANOVA (**Figure 4I**).

Collectively, these findings demonstrate that gut-specific loss of Kdm5 is sufficient to impair social behavior and that *L. plantarum* Lp39 strain produces a qualitative improvement in social proximity that does not reach statistical significance, highlighting a dissociation between probiotic effects on microbial composition and social behavior. Notably, while *L. helveticus* improves social proximity in whole-body *Kdm5^LOF^* flies, this effect is not recapitulated in the gut-specific *Kdm5* knockdown model.

Consistent with observations in the whole-body *Kdm5* loss-of-function mutant, open field testing revealed no significant differences in spatial exploration or zone occupancy across gut-specific *luciferase^RNAi^* controls or *Kdm5^RNAi^*flies under any probiotic treatment condition (**Supplementary Figure 2).**

### Fecal microbiota transplantation alters gut bacterial abundance in *Kdm5^LOF^* flies

To test whether a transfer of gut microbiota from healthy flies could restore microbial deficits observed in *Kdm5^LOF^* flies, we performed a fecal microbiota transplantation (FMT) paradigm in which adult control and *Kdm5^LOF^*flies served as both donors and recipients (**Figure 5A**). To validate the FMT paradigm and define the exposure window, we independently verified bacterial transfer using a fluorescently labeled *Lactiplantibacillus plantarum* strain constitutively expressing mCherry. Using this approach, we confirmed that donor-derived bacteria were detectable in recipient fly guts following a 3-day donor conditioning period and 24-h recipient exposure (**Supplementary Fig. 3**), supporting the efficacy of the FMT protocol used in subsequent experiments.

**Fig. 5.**
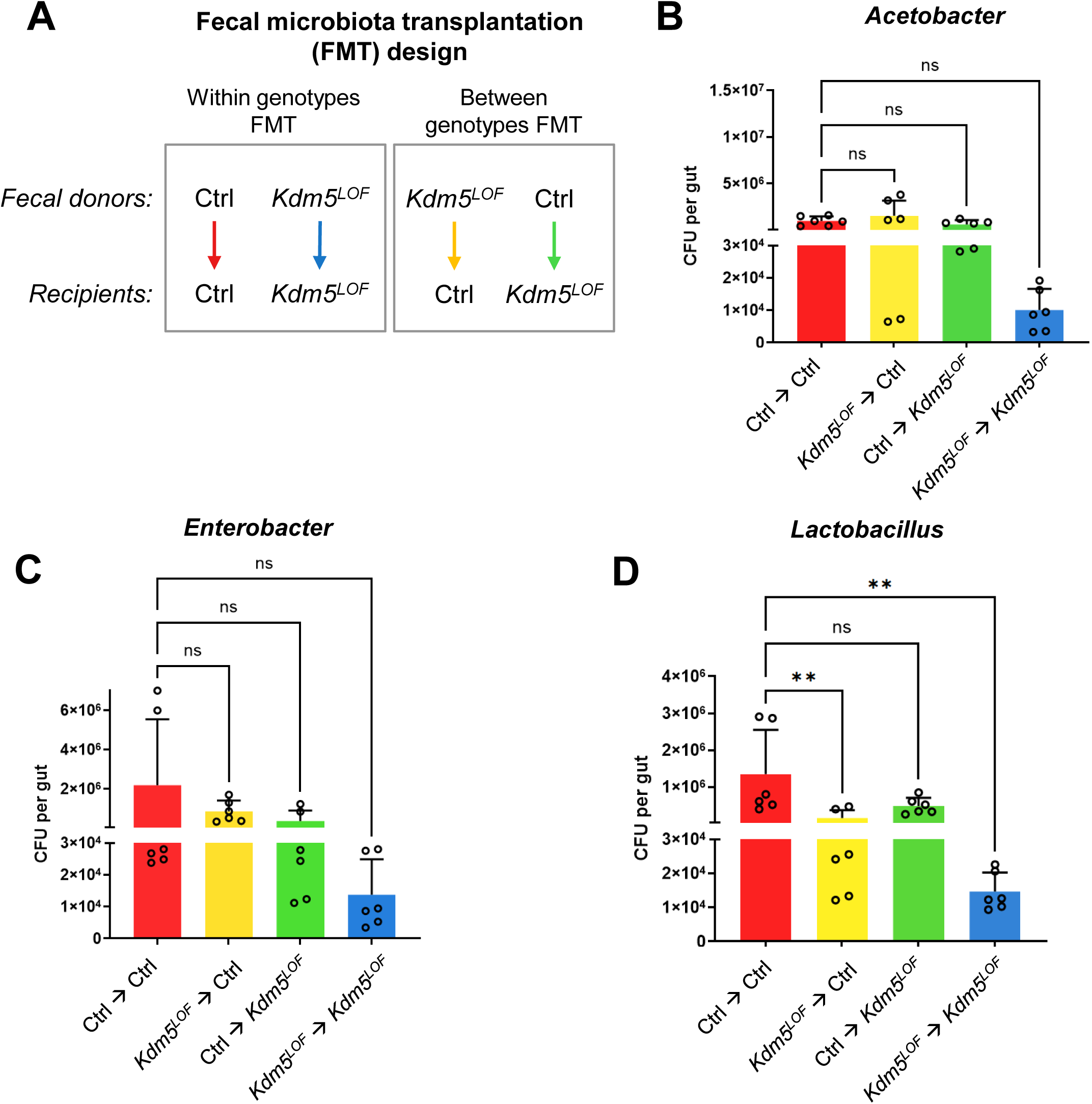
Fecal microbiota transplantation selectively restores intestinal bacterial abundance in *Kdm5^LOF^*flies. (A) Schematic of the fecal microbiota transplantation (FMT) experimental design. Young adult control and *Kdm5^LOF^* flies (5–7 days old) were used as fecal donors and maintained on standard food for 3 days. Donor flies were then removed, and age-matched recipient flies were transferred into the conditioned vials and exposed to donor fecal material for 24 h. Donor–recipient combinations are indicated. (B–D) Following FMT, guts were dissected from recipient flies, homogenized in sterile PBS, serially diluted, and plated on selective agar media to quantify CFUs of *Acetobacter* (B), *Enterobacter* (C), and *Lactobacillus* spp. (D). Bacterial colonies were counted after 24–48 h of incubation, and CFUs were calculated per gut. Bars represent mean ± SEM from three independent biological replicates, with duplicate technical platings per sample. Statistical significance was determined using one-way ANOVA followed by Dunnett’s multiple-comparisons test, as indicated. **p < 0.01; ns, not significant.

Recipient flies were exposed for 24 h to food previously conditioned with fecal material from either control or *Kdm5^LOF^* donor flies, allowing ingestion of donor-associated microbiota. Gut bacterial abundance was subsequently assessed by CFU analysis.

CFU quantification revealed no statistically significant differences in *Acetobacter* abundance among the four donor–recipient combinations (**Figure 5B**). Although *Kdm5^LOF^* recipient flies exposed to fecal material from *Kdm5^LOF^* donors showed a trend toward reduced *Acetobacter* levels compared to other groups, this effect did not reach statistical significance. Similarly, exposure of *Kdm5^LOF^*recipients to control-derived fecal material resulted in a modest increase in *Acetobacter* abundance that was not statistically significant. Control recipient flies exhibited stable *Acetobacter* levels regardless of donor genotype.

A similar pattern was observed for *Enterobacter* abundance (**Figure 5C**). Control recipient flies showed no significant changes in *Enterobacter* levels following exposure to either control or *Kdm5^LOF^* donor fecal material. *Kdm5^LOF^* recipient flies exposed to *Kdm5^LOF^*donor feces exhibited reduced *Enterobacter* abundance, whereas exposure to control donor feces produced a modest, non-significant increase.

In contrast, FMT produced pronounced effects on *Lactobacillus* abundance (**Figure 5D**). Control recipient flies exposed to control donor fecal material maintained high *Lactobacillus* levels, whereas exposure to fecal material from *Kdm5^LOF^*donors resulted in a significant reduction in *Lactobacillus* abundance. Notably, *Kdm5^LOF^* recipient flies exhibited significantly reduced *Lactobacillus* levels when exposed to *Kdm5^LOF^* donor fecal material; however, transplantation of fecal material from control donors significantly increased *Lactobacillus* abundance in *Kdm5^LOF^* recipients, restoring levels toward those observed in control flies.

Collectively, these findings indicate that fecal microbiota transplantation selectively influences gut bacterial composition in a genotype-dependent manner and that transfer of microbiota from healthy donors is sufficient to restore *Lactobacillus* abundance in *Kdm5^LOF^* flies.

### FMT induces donor-dependent shifts in gut microbial diversity and community structure

To determine whether FMT alters gut microbial communities in recipient flies, we performed full-length 16S rRNA gene sequencing on control and *Kdm5^LOF^* recipient flies following exposure to fecal material from control or *Kdm5^LOF^*donors (**Figure 6**).

**Fig. 6.**
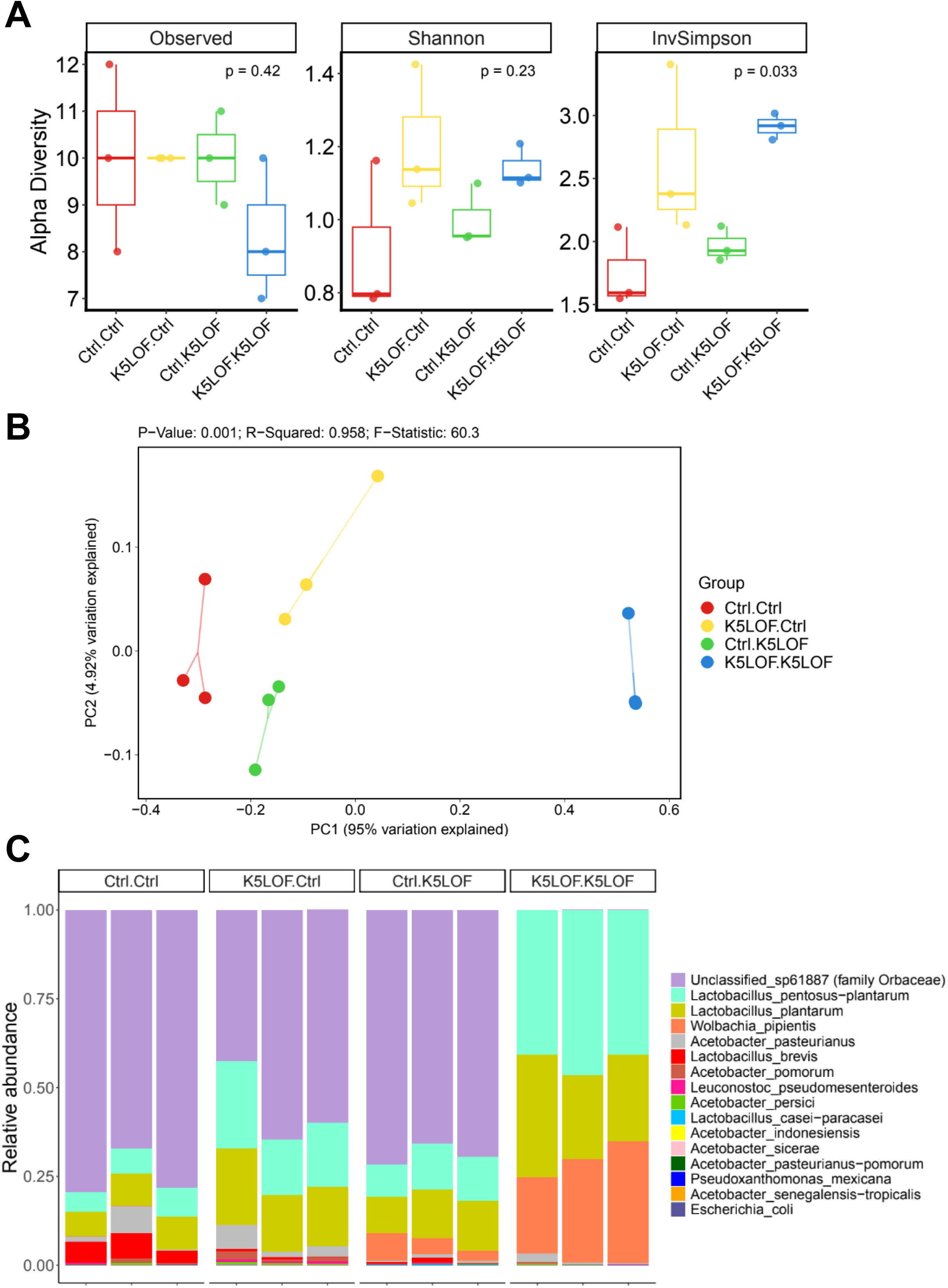
FMT induces donor-dependent remodeling of the gut microbiome in *Kdm5^LOF^* flies. FMT was performed as described in Figure 5. Briefly, young adult control and *Kdm5^LOF^* flies (5–7 days old) were exposed for 24 h to fecal material derived from either control or *Kdm5^LOF^* donor flies prior to gut dissection and microbiome analysis. (A) Alpha diversity metrics, including observed species, Shannon diversity, and inverse Simpson diversity, were calculated for recipient flies following FMT. (B) Principal coordinates analysis (PCoA) of Bray–Curtis dissimilarities showing separation of gut microbial communities based on donor–recipient combinations. PERMANOVA statistics are indicated. (C) Relative abundance of dominant bacterial taxa at the species level in recipient flies following FMT. Each condition represents three independent biological replicates, with ten pooled guts per replicate. Statistical analyses for alpha diversity were performed as indicated; community-level differences were assessed by PERMANOVA.

Alpha diversity analyses revealed modest effects of FMT on overall community richness and evenness (**Figure 6A**). No significant differences were observed in observed species or Shannon diversity across donor–recipient combinations. In contrast, the Inverse Simpson Index differed significantly among groups (Kruskal-Wallis; p = 0.033), indicating changes in community dominance structure following FMT. Notably, *Kdm5^LOF^*recipient flies exposed to *Kdm5^LOF^* donor microbiota exhibited increased inverse Simpson diversity relative to *Kdm5^LOF^* recipients exposed to control donor microbiota, suggesting differences in community eveness and dominance rather than changes in overall richness.

PCoA of Bray–Curtis dissimilarities demonstrated clear separation of gut microbial communities based on donor–recipient combinations (**Figure 6B**). PERMANOVA analysis confirmed that community composition differed significantly among groups (p = 0.001; R² = 0.958), indicating that both donor and recipient genotypes strongly influenced microbial community structure. *Kdm5^LOF^*recipients receiving control donor microbiota clustered separately from *Kdm5^LOF^*→*Kdm5^LOF^* recipients and showed partial overlap with control-associated samples, consistent with a donor-dependent restructuring of the gut microbiome.

Analysis of relative taxonomic abundance revealed genotype-dependent differences in dominant bacterial taxa following FMT (**Figure 6C**). Control recipient flies exhibited microbiota enriched by *Lactobacillus* and *Acetobacter* species. In contrast, *Kdm5^LOF^*→*Kdm5^LOF^* recipients showed a microbiota characterized by high relative abundance of *Wolbachia pipientis* together with substantial representation of *Lactobacillus* taxa, including *Lactobacillus pentosus–plantarum* and *Lactobacillus plantarum*. Although *Lactobacillus* species were detectable across all FMT groups, their relative contribution to overall community composition varied depending on donor–recipient genotype combinations.

### Differential abundance analysis identifies *Lactobacillus* taxa associated with *Kdm5^LOF^* recipient microbiomes following FMT

To identify specific bacterial taxa associated with FMT–mediated remodeling of the gut microbiome, we performed differential abundance analysis on the full-length 16S rRNA gene sequencing data from recipient flies (**Figure 7**).

**Fig. 7.**
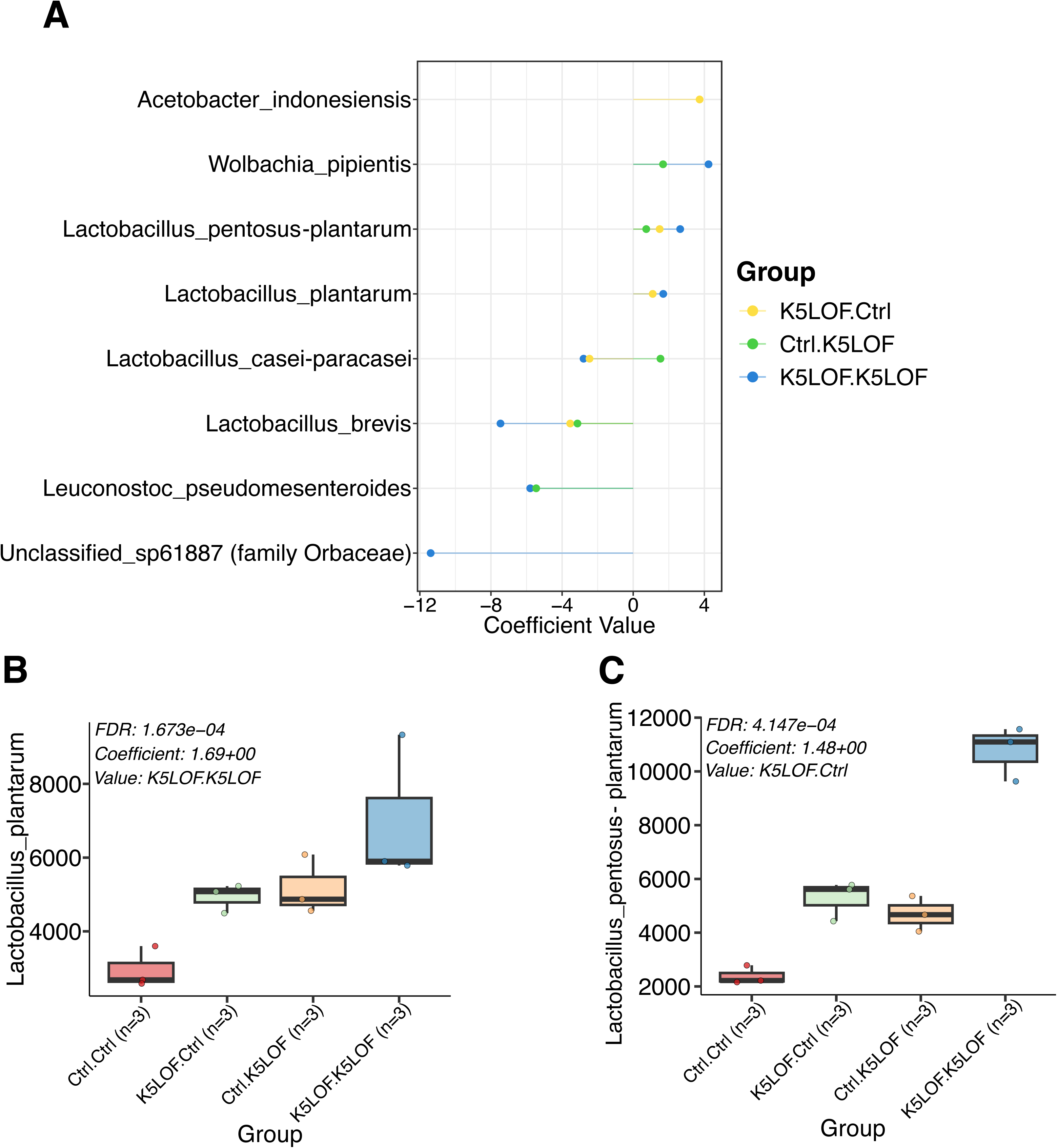
Differential abundance analysis identifies enrichment of *Lactobacillus* taxa following FMT. Differential abundance analysis was performed on 16S rRNA gene sequencing data from recipient flies following FMT. (A) Coefficient plot showing differential abundance of selected bacterial taxa across donor–recipient combinations following FMT. Positive coefficient values indicate relative enrichment, whereas negative values indicate relative depletion. (B–C) Relative abundance of *Lactobacillus plantarum* (B) and *Lactobacillus pentosus–plantarum* (C) in control and *Kdm5^LOF^* recipient flies following FMT. Individual points represent biological replicates. Each condition represents three independent biological replicates. Statistical significance was determined using differential abundance analysis with false discovery rate (FDR) correction. Adjusted p-values and coefficients are indicated where significant.

Differential abundance analysis revealed donor-dependent shifts in several bacterial taxa across FMT conditions (**Figure 7A**). Among these, *Lactobacillus* species showed the most consistent changes. In particular, *Lactobacillus pentosus–plantarum* and *Lactobacillus plantarum* showed significant positive associations with the *Kdm5^LOF^*→*Kdm5^LOF^*condition, indicating that these taxa were statistically enriched in *Kdm5^LOF^* recipients exposed to *Kdm5^LOF^*donor microbiota. In contrast, taxa associated with *Wolbachia* were also enriched in the *Kdm5^LOF^*→*Kdm5^LOF^* condition, consistent with increased dominance of *Wolbachia* observed in relative abundance analyses (**Figure 6C**).

Consistent with the differential abundance results, relative abundance plots demonstrated that *Lactobacillus plantarum* and *Lactobacillus pentosus–plantarum* reached their highest levels in *Kdm5^LOF^*→*Kdm5^LOF^* recipients, with Ctrl→*Kdm5^LOF^* recipients exhibiting intermediate abundances (**Figure 7B,C**). These taxa were present across all groups, but their relative enrichment and contribution to microbiome structure differed depending on donor–recipient genotype combinations.

Collectively, these findings indicate that FMT results in taxon-specific remodeling of the gut microbiome in *Kdm5^LOF^* flies and that *Lactobacillus* taxa are key contributors to genotype-associated microbial community states following transplantation.

### FMT partially rescues social behavior deficits in *Kdm5^LOF^* flies

To assess whether FMT influences social behavior, we performed the social distance assay on recipient flies following exposure to fecal material from control or *Kdm5^LOF^* donors using the same experimental paradigm described above (**Figure 8**). Control recipient flies receiving fecal microbiota from either control or *Kdm5^LOF^* donors exhibited normal social behavior, characterized by a high proportion of flies occupying the first distance bin, indicative of close social spacing (**Figure 8A, B**).

**Fig. 8.**
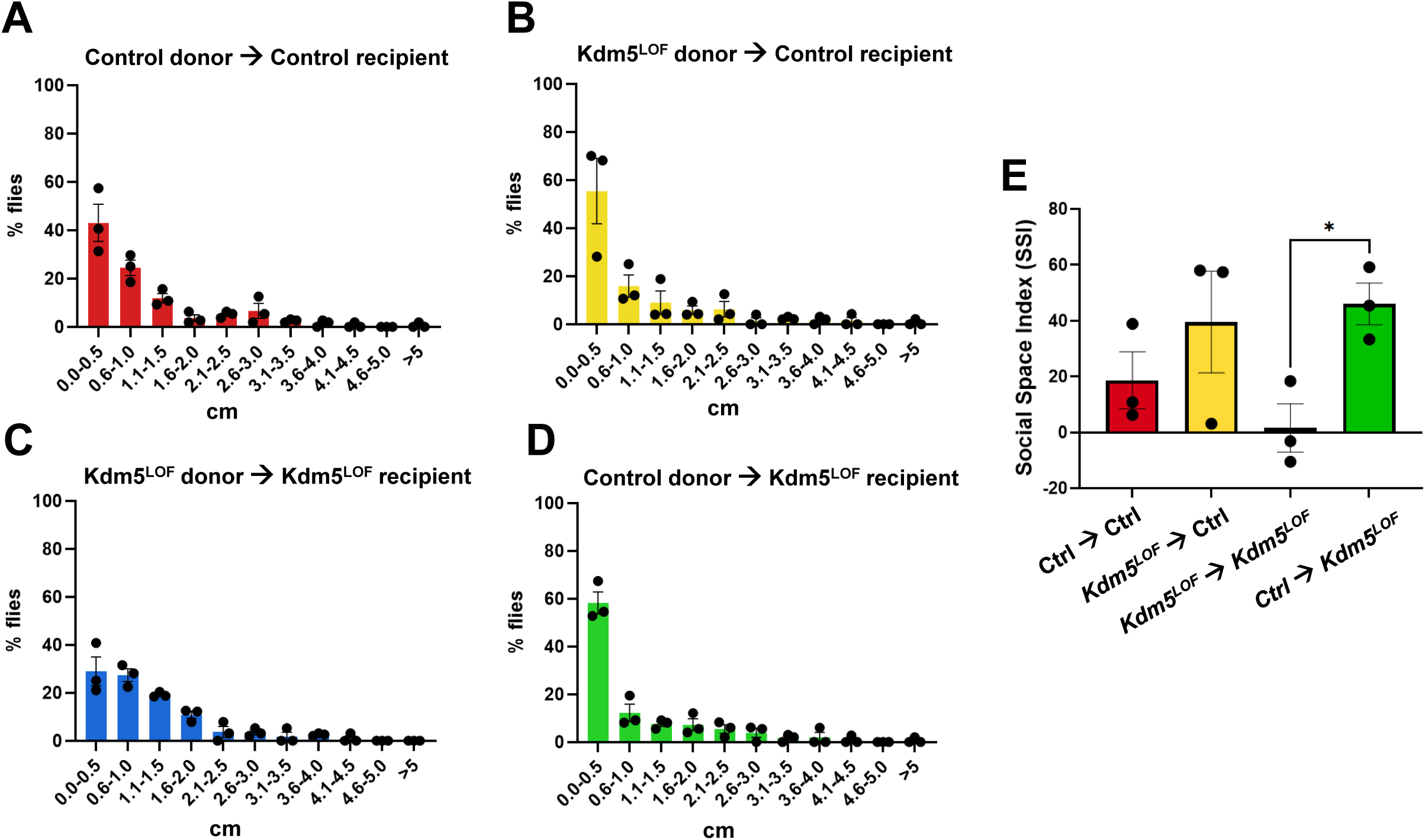
FMT partially rescues social behavior deficits in *Kdm5^LOF^* flies. Young adult control and *Kdm5^LOF^* flies (5–7 days old) were subjected to FMT as described in Figure 5. Briefly, recipient flies were exposed for 24 hours to fecal material derived from either control or *Kdm5^LOF^* donor flies prior to behavioral testing. Social spacing behavior was assessed using the Social Distance assay. Histograms (A–D) show the percentage of flies occupying 0.5-cm distance bins, calculated based on the distance from the nearest to the farthest neighbor. Panels correspond to the following donor–recipient combinations: control→control (A), Kdm5LOF→control (B), Kdm5LOF→Kdm5LOF (C), and control→Kdm5LOF (D). (E) The Social Space Index (SSI) was calculated as the percentage of flies in the first distance bin minus the percentage in the second bin for each condition. Positive SSI values indicate increased social proximity, whereas values at or below zero indicate reduced social interaction. Bars represent mean ± SEM from three independent biological replicates. Statistical analysis was performed using one-way ANOVA. A planned pairwise comparison between *Kdm5^LOF^*→*Kdm5^LOF^*and control→*Kdm5^LOF^*F groups was conducted using Holm–Šídák’s multiple-comparisons test. *p < 0.05.

In contrast, *Kdm5^LOF^* recipient flies exposed to fecal microbiota from *Kdm5^LOF^* donors displayed impaired social behavior, with a shift toward increased occupancy of the second distance bin, consistent with reduced social interaction (**Figure 8C**). Notably, *Kdm5^LOF^*recipient flies receiving fecal microbiota from control donors (control→*Kdm5^LOF^*) exhibited a redistribution toward shorter inter-fly distances relative to *Kdm5^LOF^*→*Kdm5^LOF^*recipients (**Figure 8D**).

Quantification of social behavior using the Social Space Index (SSI) revealed that although a one-way ANOVA did not detect a significant overall effect of donor–recipient group on SSI (p = 0.10), a planned pairwise comparison demonstrated that *Kdm5^LOF^*recipient flies receiving control donor microbiota exhibited significantly higher SSI values than *Kdm5^LOF^*recipients receiving *Kdm5^LOF^* donor microbiota (Holm–Šídák–adjusted p = 0.0297; **Figure 8E**). Together, these results indicate that fecal microbiota from healthy donors partially rescues social interaction deficits in *Kdm5^LOF^* flies.

## Discussion

In this study, we demonstrate that loss of the chromatin regulator *Kdm5* disrupts gut microbial homeostasis and social behavior in *Drosophila melanogaster*, and that these phenotypes can be differentially modulated by probiotic supplementation and fecal microbiota transplantation. Across multiple experimental paradigms, including whole-body *Kdm5* loss-of-function mutants, gut-specific *Kdm5* knockdown, and FMT, we consistently observed reduced abundance of culturable *Lactobacillus* species, supporting a central role for *Kdm5* in maintaining a gut environment permissive for beneficial microbial colonization.

Consistent with previous work by Chen et al. (2019) [36], *Kdm5^LOF^*mutant flies exhibited gut microbial dysbiosis characterized by reduced *Lactobacillus* abundance. Extending these findings, we show that dysbiosis in *Kdm5^LOF^* mutants is not limited to *Lactobacillus*, but also includes reductions in *Acetobacter* and *Enterobacter* CFUs. Supplementation with *Lactiplantibacillus plantarum* Lp39 robustly restored the abundance of all three genera, whereas *Lactobacillus helveticus* had minimal effects on microbial composition. These results suggest that *L. plantarum* plays a foundational role in shaping the *Drosophila* gut microbiota, potentially by facilitating colonization or growth of other commensal taxa, consistent with recent reports identifying *L. plantarum* as an early colonizer that promotes *Acetobacter* establishment in the adult gut [48–50].

Notably, restoration of gut microbial abundance was not sufficient to rescue social behavior defects. While *L. plantarum* effectively corrected microbial dysbiosis in *Kdm5^LOF^* mutant flies, it produced little to no improvement in social behavior. In contrast, *L. helveticus* significantly rescued social interaction deficits despite having minimal impact on overall gut bacterial abundance. This dissociation between microbial abundance and behavioral outcomes indicates that probiotic-mediated behavioral rescue depends on functional properties of specific bacterial strains rather than bulk microbial restoration alone. Such properties may include production of neuroactive metabolites, modulation of host signaling pathways, or immune interactions independent of bacterial load (reviewed in [7,51,52]).

Strain-specific effects were further highlighted by differences between *L. plantarum* isolates. The Lp39 strain used in this study, originally isolated from fermented cabbage, differs from the L168 strain used by Chen et al. (2019) [36], which was isolated from the guts of young wild-type female *Drosophila*. Prior work has shown that different *L. plantarum* strains vary widely in their capacity to produce neurotransmitters such as GABA, histamine, and acetylcholine (reviewed in [51]). Whether such strain-specific metabolic outputs contribute to behavioral rescue in *Kdm5* mutants remains an important question for future investigation. Similarly, the ability of *L. helveticus* to rescue social behavior in *Kdm5^LOF^*flies despite limited effects on microbial abundance raises the possibility that this strain influences host behavior through metabolite production or signaling mechanisms independent of stable gut colonization. Consistent with this idea, studies in humans, mice, and rats have shown that *L. helveticus* can modulate host behavior, including reductions in anxiety and improvements in cognitive function, although in some cases *L. helveticus* was administered in combination with other probiotics or as part of a fermented product [53–56]. Further studies are needed to define the specific molecular mechanisms by which *L. helveticus* influences gut–brain communication.

Our gut-specific *Kdm5* knockdown experiments further demonstrate that intestinal loss of *Kdm5* is sufficient to induce microbial dysbiosis and social behavior deficits, implicating the gut epithelium as a critical site of *Kdm5* function. Although probiotic supplementation restored microbial abundance in this model, behavioral rescue was modest, reinforcing the notion that microbial composition and behavioral outcomes can be uncoupled. Differences observed between whole-body mutants and gut-specific knockdown controls also suggest that host genetic background influences susceptibility to probiotic-induced microbial and behavioral changes. These findings indicate that *Kdm5* function within the gut epithelium is sufficient to influence microbial viability and social behavior, independent of broader developmental effects.

Fecal microbiota transplantation provided additional evidence linking gut microbial state to social behavior. Transfer of microbiota from healthy donors selectively restored *Lactobacillus spp.* abundance and partially rescued social interaction deficits in *Kdm5* recipient flies. In contrast, fecal material from *Kdm5* donors significantly reduced *Lactobacillus* CFUs in control recipients without affecting behavior, suggesting that dysbiotic microbiota can alter microbial community structure in otherwise healthy hosts without producing overt behavioral consequences. These findings highlight the context-dependent effects of microbiota transfer and underscore the complexity of host–microbe–behavior interactions. Notably, they parallel emerging clinical and preclinical studies reporting behavioral improvements following microbiota transfer in subsets of individuals with ASD while also emphasizing that microbiome-mediated effects on behavior are partial, host-dependent, and unlikely to be universally predictive (reviewed in [57]).

Across CFU-based and full-lenght 16S rRNA sequencing analyses, we observed an apparent discrepancy whereby *Lactobacillus* taxa remained detectable by sequencing despite reduced culturability in *Kdm5*-deficient flies. This likely reflects fundamental methodological differences: CFU assays quantify viable, actively replicating bacteria under defined growth conditions, whereas sequencing captures relative taxonomic representation independent of metabolic state. The persistence of *Lactobacillus* DNA alongside reduced CFUs suggests that loss of *Kdm5* alters the gut environment in a manner that impairs bacterial viability or growth rather than eliminating these taxa entirely. Increased dominance of the insect endosymbiont *Wolbachia pipientis* [58] in *Kdm5*-deficient microbiomes may further contribute to reduced community evenness and competitive exclusion of metabolically active gut bacteria. Consistent with this possibility, *Wolbachia*-infected adult *Drosophila* (wMel strain) have been shown to exhibit reduced abundance of *Acetobacter* ssp. including *Acetobacter pasteurianus* [59], a phenotype also observed in our 16S rRNA sequencing data. Importantly, how an obligate intracellular endosymbiont such as *Wolbachia* influences microbial populations residing in the gut lumen remains unresolved, and may involve indirect effects on host physiology, immune signaling, or metabolic environment rather than direct microbial competition. Together, these findings indicate that *Kdm5* plays a critical role in maintaining a gut environment permissive for *Lactobacillus* viability and growth, a phenotype conserved across genetic and microbiota-transfer contexts

Importantly, microbial dysbiosis in *Kdm5* mutants occurred in the absence of detectable intestinal barrier dysfunction. Neither survival nor gut permeability differed across genotypes or treatments in the Smurf assay for the timeframe studied, indicating that microbial changes are not secondary to overt epithelial breakdown. This contrasts with interpretations of earlier dye-based assays and underscores the value of established barrier integrity measures when assessing gut dysfunction.

In conclusion, this study identifies *Kdm5* as a critical regulator of gut microbial homeostasis and social behavior in *Drosophila melanogaster*. Our findings demonstrate that loss of *Kdm5* disrupts the viability and balance of core gut bacterial taxa and leads to impairments in social interaction, effects that can be differentially modulated by probiotic supplementation and fecal microbiota transplantation. Notably, probiotic and microbiota-based interventions exert strain-specific and phenotype-specific effects, underscoring the complexity of host–microbe–behavior interactions and revealing a dissociation between microbial abundance and behavioral rescue. Rather than simply shaping which microbes are present, *Kdm5* appears to govern whether beneficial microbes can thrive and exert functional effects on host behavior. Together, these results provide a framework for dissecting how chromatin regulators influence gut–brain communication and offer insight into microbiota-mediated modulation of social behavior relevant to neurodevelopmental disorders.

### Limitations and future directions

Several limitations of this study warrant consideration and point toward important future directions. First, although probiotic and FMT-mediated effects on social behavior were reproducible across experiments, behavioral assays are inherently variable, and increasing sample size may provide greater statistical power to detect additional or more subtle effects, particularly in gut-specific *Kdm5* knockdown models. Second, the temporal stability of probiotic- and FMT-induced changes remains unknown. Whether microbial and behavioral rescue effects persist long-term or require continuous exposure will be important to determine in future longitudinal studies.

Third, the mechanism by which *Lactobacillus helveticus* rescues social behavior despite minimal effects on culturable gut bacteria remains unresolved. This suggests that behavioral modulation may occur through microbial metabolites, indirect host signaling, or interactions with microbial community members not captured by CFU-based analyses. In this context, future studies incorporating metabolomics or host transcriptomic approaches will be valuable.

In addition, our analyses focused primarily on bacterial taxa identifiable using full-length 16S rRNA gene sequencing, yet other components of the gut microbiome, including viruses and fungi, may also contribute to host–microbe–behavior interactions. Indeed, a study in individuals with ASD reported alterations not only in the Bacillota (formerly Firmicutes) to Bacteroidetes ratio but also in the relative abundance of the fungal genus *Candida* [5]. Comprehensive metagenomic or multi-omics approaches will be necessary to capture these additional layers of complexity, which will also provide information on potential associated metabolites, genes, pathways and enzymes. Finally, all experiments in this study were performed on female flies, chosen due to their robust intestinal regenerative capacity; whether similar microbiota–behavior relationships exist in males remains to be determined. Moreover, although our findings reveal strong associations between microbial composition and behavior, future work will be required to establish direct causal links between specific microbial factors, host pathways, and behavioral outputs. For instance, future studies directly testing the contribution of innate immune signaling downstream of Kdm5—proposed by Chen et al. (2019) [36] to involve chronic IMD/Relish pathway activation—will be necessary to determine how this immune axis interacts with strain-specific probiotic effects and to disentangle immune-dependent and -independent mechanisms of behavioral modulation.

## Acknowledgements

We thank Dr. William Ludington (Carnegie Science) for donating the LpWF-mCherry strain containing pCD256NS-mCherry, a modified version of the plasmid pCD256-mCherry originally generated by Stefan Heinl and Reingard Grabherr (University of Natural Resources and Life Sciences, BOKU, Vienna, Austria).

We are also grateful to Dr. Alfredo Ghezzi (University of Puerto Rico, Río Piedras), Dr. Alberto Cruz-Martín (University of Colorado, Anschutz Medical Campus), and Dr. Alberto Sabat (University of Puerto Rico, Río Piedras) for their valuable feedback and guidance as members of Natalia Peta’s thesis committee. Thanks to Dr. Esther Peterson (University of Puerto Rico, Río Piedras) for the use of the Nikon fluorescent microscope.

Finally, we thank Dr. Ricardo Chiesa and Geraldine M. Ortiz Sosa (University of Puerto Rico, Cayey) for their insight regarding the Open Field Test behavioral assay.

ChatGPT (OpenAI, 2024) was used to refine grammar and phrasing in parts of the manuscript. All research, analysis, and manuscript content were solely produced by the authors.

## Statements

### Statement of Ethics

This study used *Drosophila melanogaster* and did not involve human participants or vertebrate animals. As such, ethics approval was not required

### Author Contributions

N.A.P.M. was responsible for conceptualization, the majority of experimental work, data analysis, interpretation, and writing and reviewing the manuscript.

M.R.A. designed and tested the fecal microbiota transplantation (FMT) protocol and contributed to manuscript revision.

T.M.S.R. contributed to data analysis of FMT/16S experiment and bioinformatic support and contributed to manuscript revision.

I.A.R.F. contributed to conceptualization, performed the FMT/16S experiments, secured funding, provided guidance and mentorship, contributed to data interpretation and analysis, and was involved in writing and reviewing the manuscript.

### Data Availability Statement

16S sequencing data are available in the NCBI Sequence Read Archive (SRA) under BioProject accession number PRJNA1365331. Additional data generated during this study are available from the corresponding author upon reasonable request.

## References

1 Shaw K.A., Williams S, Patrick MA. Prevalence and Early Identification of Autism Spectrum Disorder Among Children Aged 4 and 8 Years - Autism and Developmental Disabilities Monitoring Network, 16 Sites, United States, 2022. MMWR Surveill Summaries. 2025 Apr;74(74):1–22.

2 American Psychiatric Association. Diagnostic and Statistical Manual of Mental Disorders. 5th ed. Washington, DC: American Psychiatric Publishing; 2022.

3 Genovese A, Butler MG. The Autism Spectrum: Behavioral, Psychiatric and Genetic Associations. Genes (Basel). 2023 Mar;14(3). DOI: 10.3390/genes14030677

4 Ibrahim SH, Voigt RG, Katusic SK, Weaver AL, Barbaresi WJ. Incidence of gastrointestinal symptoms in children with autism: A population-based study. Pediatrics. 2009 Aug;124(2):680–6.

5 Strati F, Cavalieri D, Albanese D, De Felice C, Donati C, Hayek J, et al. New evidences on the altered gut microbiota in autism spectrum disorders. Microbiome. 2017;5(1). DOI: 10.1186/s40168-017-0242-1

6 Salim S, Banu A, Alwa A, Gowda SBM, Mohammad F. The gut-microbiota-brain axis in autism: what Drosophila models can offer? J Neurodev Disord. 2021 Dec;13(1). DOI: 10.1186/s11689-021-09378-x

7 Sharon G, Sampson TR, Geschwind DH, Mazmanian SK. The Central Nervous System and the Gut Microbiome. Cell. 2016 Nov;167(4):915–32.

8 Cryan JF, O KJ, M Cowan CS, Sandhu K V, S Bastiaanssen TF, Boehme M, et al. The Microbiota-Gut-Brain Axis. Physiol Rev. 2019;99:1877–2013.

9 Morais LH, Schreiber HL, Mazmanian SK. The gut microbiota–brain axis in behaviour and brain disorders. Nat Rev Microbiol. 2021 Apr;19(4):241–55.

10 Hill C, Guarner F, Reid G, Gibson GR, Merenstein DJ, Pot B, et al. Expert consensus document: The international scientific association for probiotics and prebiotics consensus statement on the scope and appropriate use of the term probiotic. Nat Rev Gastroenterol Hepatol. 2014;11(8):506–14.

11 Buffington SA, Di Prisco GV, Auchtung TA, Ajami NJ, Petrosino JF, Costa-Mattioli M. Microbial Reconstitution Reverses Maternal Diet-Induced Social and Synaptic Deficits in Offspring. Cell. 2016 Jun;165(7):1762–75.

12 Tabouy L, Getselter D, Ziv O, Karpuj M, Tabouy T, Lukic I, et al. Dysbiosis of microbiome and probiotic treatment in a genetic model of autism spectrum disorders. Brain Behav Immun. 2018 Oct;73:310–9.

13 Hsiao EY, McBride SW, Hsien S, Sharon G, Hyde ER, McCue T, et al. Microbiota modulate behavioral and physiological abnormalities associated with neurodevelopmental disorders. Cell. 2013 Dec;155(7):1451–63.

14 Sgritta M, Dooling SW, Buffington SA, Momin EN, Francis MB, Britton RA, et al. Mechanisms Underlying Microbial-Mediated Changes in Social Behavior in Mouse Models of Autism Spectrum Disorder. Neuron. 2019 Jan;101(2):246–259.e6.

15 Mazzone L, Dooling SW, Volpe E, Uljarević M, Waters JL, Sabatini A, et al. Precision microbial intervention improves social behavior but not autism severity: A pilot double-blind randomized placebo-controlled trial. Cell Host Microbe. 2024 Jan;32(1):106–116.e6.

16 Gupta S, Allen-Vercoe E, Petrof EO. Fecal microbiota transplantation: In perspective. Therap Adv Gastroenterol. 2016 Mar;9(2):229–39.

17 Ooijevaar RE, Terveer EM, Verspaget HW, Kuijper EJ, Keller JJ. Annual Review of Medicine Clinical Application and Potential of Fecal Microbiota Transplantation. Downloaded from www.annualreviews.org Guest. 2018;70:335–51.

18 Li Y, Wang Y, Zhang T. Fecal Microbiota Transplantation in Autism Spectrum Disorder. Neuropsychiatr Dis Treat. 2022;18:2905–15.

19 Li N, Chen H, Cheng Y, Xu F, Ruan G, Ying S, et al. Fecal Microbiota Transplantation Relieves Gastrointestinal and Autism Symptoms by Improving the Gut Microbiota in an Open-Label Study. Front Cell Infect Microbiol. 2021 Oct;11. DOI: 10.3389/fcimb.2021.759435

20 Kang DW, Adams JB, Gregory AC, Borody T, Chittick L, Fasano A, et al. Microbiota Transfer Therapy alters gut ecosystem and improves gastrointestinal and autism symptoms: An open-label study. Microbiome. 2017;5(1). DOI: 10.1186/s40168-016-0225-7

21 Douglas AE. Simple animal models for microbiome research. Nat Rev Microbiol. 2019 Dec;17(12):764–75.

22 Trinder M, Daisley BA, Dube JS, Reid G. Drosophila melanogaster as a high-throughput model for host-microbiota interactions. Front Microbiol. 2017 Apr;8(APR). DOI: 10.3389/fmicb.2017.00751

23 Ludington WB, Ja WW. Drosophila as a model for the gut microbiome. PLoS Pathog. 2020 Apr;16(4). DOI: 10.1371/journal.ppat.1008398

24 Chiang MH, Ho SM, Wu HY, Lin YC, Tsai WH, Wu T, et al. Drosophila Model for Studying Gut Microbiota in Behaviors and Neurodegenerative Diseases. Biomedicines. 2022 Mar;10(3). DOI: 10.3390/biomedicines10030596

25 Buchon N, Broderick NA, Chakrabarti S, Lemaitre B. Invasive and indigenous microbiota impact intestinal stem cell activity through multiple pathways in Drosophila. Genes Dev. 2009 Oct;23(19):2333–44.

26 Broderick NA, Lemaitre B. Gut-associated microbes of Drosophila melanogaster. Gut Microbes. 2012 Jul;3(4). DOI: 10.4161/gmic.19896

27 Wong CNA, Ng P, Douglas AE. Low-diversity bacterial community in the gut of the fruitfly Drosophila melanogaster. Environ Microbiol. 2011 Jul;13(7):1889–900.

28 De Rubeis S, He X, Goldberg AP, Poultney CS, Samocha K, Cicek AE, et al. Synaptic, transcriptional and chromatin genes disrupted in autism. Nature. 2014 Nov;515(7526):209–15.

29 Satterstrom FK, Kosmicki JA, Wang J, Breen MS, De Rubeis S, An JY, et al. Large-Scale Exome Sequencing Study Implicates Both Developmental and Functional Changes in the Neurobiology of Autism. Cell. 2020 Feb;180(3):568–584.e23.

30 Liu X, Secombe J. The Histone Demethylase KDM5 Activates Gene Expression by Recognizing Chromatin Context through Its PHD Reader Motif. Cell Rep. 2015;13(10):2219–31.

31 Fieremans N, Van Esch H, de Ravel T, Van Driessche J, Belet S, Bauters M, et al. Microdeletion of the escape genes KDM5C and IQSEC2 in a girl with severe intellectual disability and autistic features. Eur J Med Genet. 2015 May;58(5):324–7.

32 Martin HC, Jones WD, Mcintyre R, Sanchez-Andrade G, Sanderson M, Stephenson JD, et al. Quantifying the contribution of recessive coding variation to developmental disorders. Science (1979). 2018 Dec;362:1–4.

33 Vallianatos CN, Farrehi C, Friez MJ, Burmeister M, Keegan CE, Iwase S. Altered gene-regulatory function of KDM5C by a novel mutation associated with autism and intellectual disability. Front Mol Neurosci. 2018 Apr;11. DOI: 10.3389/fnmol.2018.00104

34 Belalcazar HM, Hendricks EL, Zamurrad S, Liebl FLW, Secombe J. The histone demethylase KDM5 is required for synaptic structure and function at the Drosophila neuromuscular junction. Cell Rep. 2021 Feb;34(7). DOI: 10.1016/j.celrep.2021.108753

35 Drelon C, Belalcazar HM, Secombe J. The histone demethylase KDM5 is essential for larval growth in Drosophila. Genetics. 2018 Jul;209(3):773–87.

36 Chen K, Luan X, Liu Q, Wang J, Chang X, Snijders AM, et al. Drosophila Histone Demethylase KDM5 Regulates Social Behavior through Immune Control and Gut Microbiota Maintenance. Cell Host Microbe. 2019 Apr;25(4):537–552.e8.

37 Taverniti V, Guglielmetti S. Health-promoting properties of Lactobacillus helveticus. Front Microbiol. 2012;3(NOV). DOI: 10.3389/fmicb.2012.00392

38 McGuire SE, Mao Z, Davis RL. Spatiotemporal gene expression targeting with the TARGET and gene-switch systems in Drosophila. Sci STKE. 2004;2004(220). DOI: 10.1126/stke.2202004pl6

39 Micchelli CA, Perrimon N. Evidence that stem cells reside in the adult Drosophila midgut epithelium. Nature. 2006 Jan;439(7075):475–9.

40 Ohlstein B, Spradling A. The adult Drosophila posterior midgut is maintained by pluripotent stem cells. Nature. 2006 Jan;439(7075):470–4.

41 Guo L, Karpac J, Tran SL, Jasper H. PGRP-SC2 promotes gut immune homeostasis to limit commensal dysbiosis and extend lifespan. Cell. 2014;156(1–2):109–22.

42 Li H, Qi Y, Jasper H. Preventing Age-Related Decline of Gut Compartmentalization Limits Microbiota Dysbiosis and Extends Lifespan. Cell Host Microbe. 2016;19(2):240–53.

43 Simon AF, Chou MT, Salazar ED, Nicholson T, Saini N, Metchev S, et al. A simple assay to study social behavior in Drosophila: Measurement of social space within a group. Genes Brain Behav. 2012 Mar;11(2):243–52.

44 Schindelin J, Arganda-Carreras I, Frise E, Kaynig V, Longair M, Pietzsch T, et al. Fiji: An open-source platform for biological-image analysis. Nat Methods. 2012 Jul;9(7):676–82.

45 Jasper H. Annual Review of Physiology Intestinal Stem Cell Aging: Origins and Interventions. 2019 DOI: 10.1146/annurev-physiol-021119

46 Resnik-Docampo M, Sauer V, Schinaman JM, Clark RI, Walker DW, Jones DL. Keeping it tight: The relationship between bacterial dysbiosis, septate junctions, and the intestinal barrier in Drosophila. Fly (Austin). 2018 Jan;12(1):34–40.

47 Salazar AM, Resnik-Docampo M, Ulgherait M, Clark RI, Shirasu-Hiza M, Jones DL, et al. Intestinal Snakeskin Limits Microbial Dysbiosis during Aging and Promotes Longevity. iScience. 2018 Nov;9:229–43.

48 Gutiérrez-García K, Aumiller K, Dodge R, Obadia B, Deng A, Agrawal S, et al. A conserved bacterial genetic basis for commensal-host specificity. Science (1979). 2024 Dec;386:1–6.

49 Dodge R, Jones EW, Zhu H, Obadia B, Martinez DJ, Wang C, et al. A symbiotic physical niche in Drosophila melanogaster regulates stable association of a multi-species gut microbiota. Nat Commun. 2023 Dec;14(1). DOI: 10.1038/s41467-023-36942-x

50 Obadia B, Güvener ZT, Zhang V, Ceja-Navarro JA, Brodie EL, Ja WW, et al. Probabilistic Invasion Underlies Natural Gut Microbiome Stability. Current Biology. 2017 Jul;27(13):1999–2006.e8.

51 Strandwitz P. Neurotransmitter modulation by the gut microbiota. Brain Res. 2018 Aug;1693:128–33.

52 Dinan TG, Cryan JF. Gut-brain axis in 2016: Brain-gut-microbiota axis-mood, metabolism and behaviour. Nat Rev Gastroenterol Hepatol. 2017 Feb;14(2):69–70.

53 Messaoudi M, Lalonde R, Violle N, Javelot H, Desor D, Nejdi A, et al. Assessment of psychotropic-like properties of a probiotic formulation (Lactobacillus helveticus R0052 and Bifidobacterium longum R0175) in rats and human subjects. British Journal of Nutrition. 2011 Mar;105(5):755–64.

54 Alatan H, Liang S, Shimodaira Y, Wu X, Hu X, Wang T, et al. Supplementation with Lactobacillus helveticus NS8 alleviated behavioral, neural, endocrine, and microbiota abnormalities in an endogenous rat model of depression. Front Immunol. 2024;15. DOI: 10.3389/fimmu.2024.1407620

55 Chung YC, Jin HM, Cui Y, Kim DS, Jung JM, Park J Il, et al. Fermented milk of Lactobacillus helveticus IDCC3801 improves cognitive functioning during cognitive fatigue tests in healthy older adults. J Funct Foods. 2014 Sep;10:465–74.

56 Liang S, Wang T, Hu X, Luo J, Li W, Wu X, et al. Administration of Lactobacillus helveticus NS8 improves behavioral, cognitive, and biochemical aberrations caused by chronic restraint stress. Neuroscience. 2015 Dec;310:561–77.

57 Zhang J, Zhu G, Wan L, Liang Y, Liu X, Yan H, et al. Effect of fecal microbiota transplantation in children with autism spectrum disorder: A systematic review. Front Psychiatry. 2023 Mar;14:1123658.

58 Zug R, Hammerstein P. Still a host of hosts for Wolbachia: Analysis of recent data suggests that 40% of terrestrial arthropod species are infected. PLoS One. 2012 Jun;7(6). DOI: 10.1371/journal.pone.0038544

59 Simhadri RK, Fast EM, Guo R, Schultz MJ, Vaisman N, Ortiz L, et al. The Gut Commensal Microbiome of Drosophila melanogaster Is Modified by the Endosymbiont Wolbachia. mSphere. 2017 Oct;2(5). DOI: 10.1128/msphere.00287-17

