## Supplementary Materials for "Microbiota-Based Interventions Differentially Rescue Gut and Social Behavior Phenotypes in a Drosophila Autism-like Model"

### Supplementary Methods

#### Smurf Assay

Intestinal barrier integrity was assessed using the Smurf assay as previously described (Rera et al. 2011; Rera et al. 2012; Martins et al. 2018). Briefly, flies were exposed to food containing a non-absorbable blue dye, and loss of gut barrier function was scored based on systemic dye leakage. A dye solution was prepared by dissolving FD&C Blue Dye No. 1 (Erioglaucine disodium salt; Sigma-Aldrich, CAS No. 3844-45-9) in water containing 5% sucrose to a final dye concentration of 2.5% (w/v). Two hundred microliters of the dye solution were added to each food vial and thoroughly mixed to ensure uniform distribution throughout the food. Adult flies were then transferred to the dye-containing Nutri-Fly Bloomington-formula food (Genesee Scientific, Cat. No. 66-113) and maintained under standard laboratory conditions.

Flies were examined daily for two weeks using a stereomicroscope and scored as “Smurf” if blue dye was observed outside the digestive tract, indicating loss of intestinal barrier integrity. Also, “non-smurf” dead flies were also scored.

The assay was performed in three independent biological experiments, each including one technical replicate per condition containing a vial of 10-15 flies. Statistical analysis was conducted using the log-rank (Mantel–Cox) test to compare Smurf-free survival curves between groups. No statistically significant differences were detected among conditions.

#### Open Field Test

The open field test (OFT) was used to assess centrophobic (anxiety-like) behavior in flies, defined as the intrinsic tendency to avoid the center of an open arena and preferentially occupy the periphery. This assay is commonly used to evaluate changes in exploratory behavior and anxiety-like responses and can also reveal gross alterations in locomotor capacity (Soibam et al. 2012a, 2012b).

The open field arena used in this study was developed in consultation with Dr. Ricardo Chiesa (University of Puerto Rico–Cayey), who assisted in the design of the initial prototype based on Mejias et al. 2016. The arena consisted of the lid of a 14-cm-diameter Petri dish subdivided into three concentric zones representing different levels of perceived exposure: Zone 1 (center; high exposure), defined as a 2-cm-diameter central circle; Zone 2 (intermediate exposure), defined as an annular region extending 4 cm outward from the edge of Zone 1; and Zone 3 (periphery; low exposure), defined as the outermost 2-cm-wide region adjacent to the dish walls. Together, these zones encompassed the full 14-cm diameter of the arena. Zone boundaries were marked using acrylic paint, and the inner walls of the dish lid were darkened to create a visually sheltered peripheral region. The arena was placed on an illuminated background to enhance contrast during imaging.

For each trial, 6–12 flies per condition were briefly cold-anesthetized by placing vials at -20 °C for 1.5-2 mins. Anesthetized flies were positioned in the center of the arena, and a transparent plastic sheet was placed on top to prevent escape while maintaining a consistent vertical constraint. Flies were allowed to acclimate for 10 min before imaging. The arena was then photographed from above against a white background once per minute for 10 min using an iPhone smartphone (model 11).

Fly position within the arena was quantified using ImageJ/FIJI by assigning each individual to one of the three predefined zones (Zones 1–3) at each time point. Statistical analyses were performed in GraphPad Prism using a chi-square test to compare the distribution of flies across zones between genotypes and treatment conditions. For experiments testing *Kdm5*$^{LOF}$ vs control the experiments was performed five independent times (five biological replicates), with one arena per condition per replicate. For the experiments testing gut-specific manipulation of Kdm5 the experiments were performed three independent times (biological replicates).

**Verification of fecal microbiota transplantation**

To visualize bacterial transfer between donor and recipient flies during fecal microbiota transplantation (FMT), a fluorescently labeled *Lactiplantibacillus plantarum* strain was used. Specifically, the *L. plantarum* LPWF strain carrying the plasmid pCD256-mCherry-ΔEc, which constitutively expresses mCherry and carries a Chloramphenicol resistance gene, was employed to enable fluorescence-based localization of bacteria. The LPWF strain was a gift from William Ludington (Carnegie Science, John Hopkins University), and the original pCD256-mCherry plasmid was a gift from Stefan Heinl and Reinhard Grabherr (University of Natural Resources and Life Sciences, Vienna, Austria).

*L. plantarum-mCherry* was cultured in 3 mL of MRS broth (BD Difco™ Lactobacilli MRS Broth, BD288130) supplemented with 10 µg/mL chloramphenicol for 24 h at 37 °C with shaking at 200 rpm. The following day, conventionally reared, mated wild-type Canton-S female flies (>4 days old) were fasted for 3 h prior to bacterial exposure. Bacterial cultures were pelleted by centrifugation (4,000 rpm for 10 minutes at 25°C) and resuspended in 1.5 mL of 5% sucrose. Fifty microliters of the bacterial suspension were added to narrow fly food vials (Genesee Scientific, Cat. No. 32-116SB) containing approximately 10 mL of Nutri-Fly Bloomington-formula food (Genesee Scientific, Cat. No. 66-113). Donor flies were allowed to feed on the bacteria for 1–2 h.

Following bacterial feeding, donor flies were transferred to fresh vials containing Nutri-Fly Bloomington-formula food without additional bacteria and maintained for 3 days to allow deposition of fecal material. Donor flies were then removed, and age-matched recipient flies of the same genotype were introduced into the conditioned vials and allowed to feed on fecal material and food for 24 h. Recipient flies were subsequently dissected, and guts were fixed in 4% formaldehyde, washed in wash buffer (1× PBS, 0.5% bovine serum albumin and 0.1% Triton X-100) for 30 minutes and mounted using Molecular Probes™ ProLong™ Diamond Antifade Mountant with DAPI. Fluorescence imaging was performed using a Nikon E80i fluorescence microscope at 20× magnification.

### Supplementary Table 1

**Table 1.** Genotypes and Treatment Groups Used in This Study

| ID used in Figures | Genotype | Treatment |
|---|---|---|
| **Figure 1 and Figure 2** | | |
| 1 - Control, 5% suc | *$w^{1118}$; $cn^{1}$/+* | 5% sucrose (suc) |
| 2 - Control, Lp | *$w^{1118}$; $cn^{1}$/+* | *L. plantarum* (Lp) |
| 3 - Control, Lh | *$w^{1118}$; $cn^{1}$/+* | *L. helveticus* (Lh) |
| 4 - Control, Lp + Lh | *$w^{1118}$; $cn^{1}$/+* | *L. plantarum* + *L. helveticus* (Lp + Lh) |
| 5 - *Kdm5$^{LOF}$*, 5% suc | *$w^{1118}$; $kdm5^{k6801}$/$kdm5^{10424}$,$cn^{1}$* | 5% sucrose (suc) |
| 6 - *Kdm5$^{LOF}$, Lp* | *$w^{1118}$; $kdm5^{k6801}$/$kdm5^{10424}$,$cn^{1}$* | *L. plantarum* (Lp) |
| 7 - *Kdm5$^{LOF}$, Lh* | *$w^{1118}$; $kdm5^{k6801}$/$kdm5^{10424}$,$cn^{1}$* | *L. helveticus* (Lh) |
| 8 - *Kdm5$^{LOF}$, Lh + Lp* | *$w^{1118}$; $kdm5^{k6801}$/$kdm5^{10424}$,$cn^{1}$* | *L. plantarum* + *L. helveticus* (Lp + Lh) |
| **Figure 3 and Figure 4** | | |
| 1 - Control, 5% suc | *$y^{1}v^{1}$* / w*; *Myo31DF-Gal4/+; tub-Gal80$^{ts}$/UAS-lucferase$^{RNAi}$* | 5% sucrose (suc) |
| 2 - Control, Lp | *$y^{1}v^{1}$* / w*; *Myo31DF-Gal4/+; tub-Gal80$^{ts}$/UAS-lucferase$^{RNAi}$* | *L. plantarum* (Lp) |
| 3 - Control, Lh | *$y^{1}v^{1}$* / w*; *Myo31DF-Gal4/+; tub-Gal80$^{ts}$/UAS-lucferase$^{RNAi}$* | *L. helveticus* (Lh) |
| 4 - Control, Lp + Lh | *$y^{1}v^{1}$* / w*; *Myo31DF-Gal4/+; tub-Gal80$^{ts}$/UAS-lucferase$^{RNAi}$* | *L. plantarum* + *L. helveticus* (Lp + Lh) |
| 5 - *Kdm5$^{LOF}$*, 5% suc | *$y^{1}v^{1}$* / w*; *Myo31DF-Gal4/+; tub-Gal80$^{ts}$/UAS-Kdm5$^{RNAi}$* | 5% sucrose (suc) |
| 6 - *Kdm5$^{LOF}$, Lp* | *$y^{1}v^{1}$* / w*; *Myo31DF-Gal4/+; tub-Gal80$^{ts}$/UAS-Kdm5$^{RNAi}$* | *L. plantarum* (Lp) |
| 7 - *Kdm5$^{LOF}$, Lh* | *$y^{1}v^{1}$* / w*; *Myo31DF-Gal4/+; tub-Gal80$^{ts}$/UAS-Kdm5$^{RNAi}$* | *L. helveticus* (Lh) |
| 8 - *Kdm5$^{LOF}$, Lh + Lp* | *$y^{1}v^{1}$* / w*; *Myo31DF-Gal4/+; tub-Gal80$^{ts}$/UAS-Kdm5$^{RNAi}$* | *L. plantarum* + *L. helveticus* (Lp + Lh) |

| **Figure 5-8** | | |
|---|---|---|
| **ID used in Figures** | **Genotype (Recipient)** | **Treatment (Fecal Donor Genotype)** |
| Ctrl → Ctrl | $w^{1118}$; $cn^{1}/+$ | Feces from $w^{1118}$; $cn^{1}/+$ |
| $Kdm5^{LOF}$→ $Kdm5^{LOF}$ | $w^{1118}$; $kdm5^{k6801}/kdm5^{10424},cn^{1}$ | Feces from $w^{1118}$; $kdm5^{k6801}/kdm5^{10424},cn^{1}$ |
| $Kdm5^{LOF}$→ Ctrl | $w^{1118}$; $kdm5^{k6801}/kdm5^{10424},cn^{1}$ | Feces from $w^{1118}$; $kdm5^{k6801}/kdm5^{10424},cn^{1}$ |
| Ctrl → $Kdm5^{LOF}$ | $w^{1118}$; $cn^{1}/+$ | Feces from $w^{1118}$; $cn^{1}/+$ |

### Supplementary Results

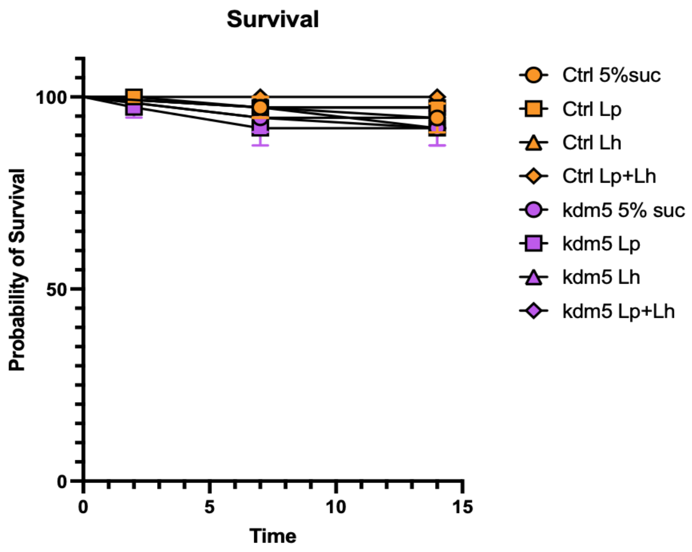


**Supplementary Fig. 1**. **Survival of control and *Kdm5*$^{LOF}$ flies following probiotic treatment.** Adult female flies were maintained on Smurf food supplemented with 5% sucrose (mock), *Lactiplantibacillus plantarum* (Lp), *Lactobacillus helveticus* (Lh), or a 1:1 combination of Lp and Lh for 14 days. Survival probability was monitored over time. Data represent the mean ± SEM from three independent biological experiments per condition. Statistical analysis was performed using the log-rank (Mantel–Cox) test to compare survival curves; no significant differences were detected among groups. No Smurf phenotype was observed during the monitoring period (0/37 flies per group).

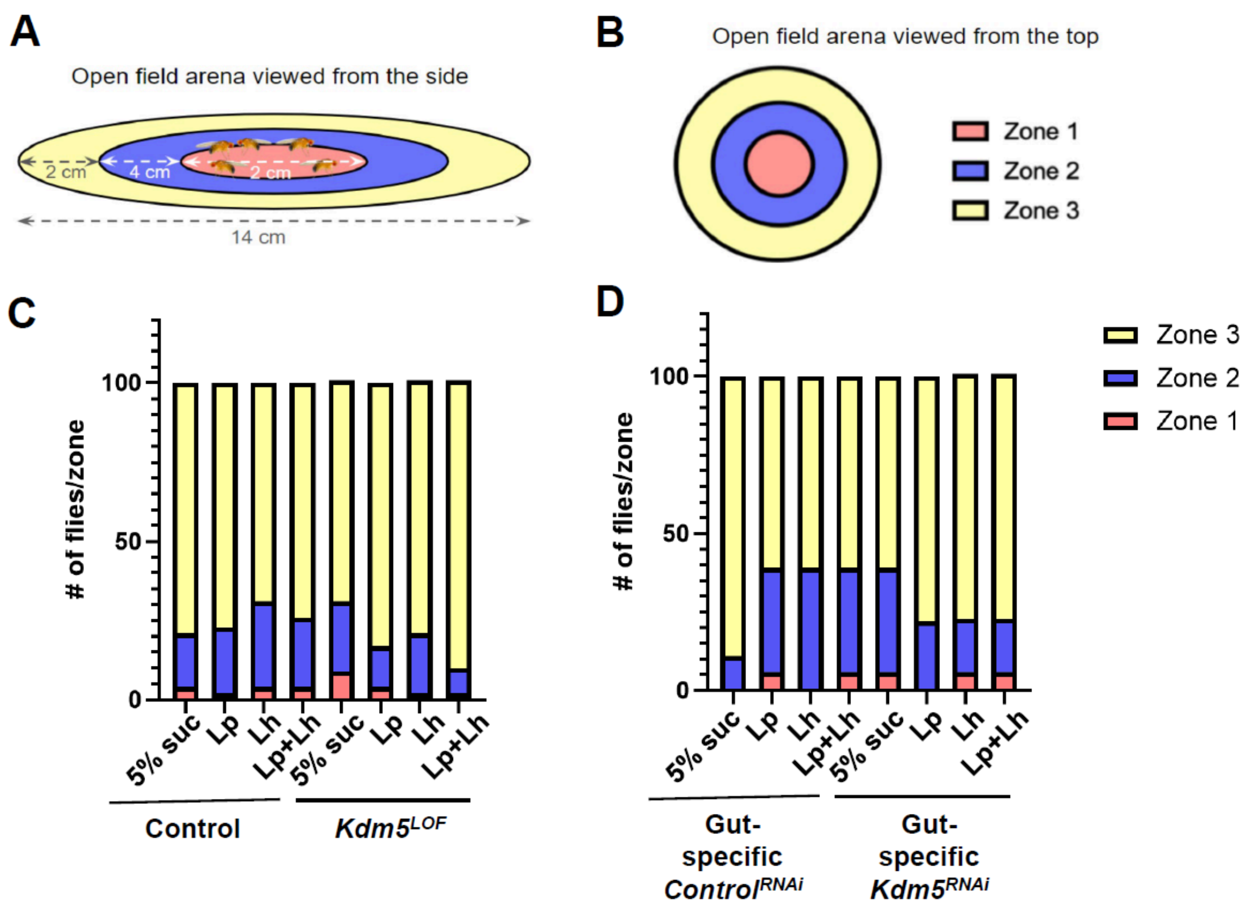


**Supplementary Fig. 2. Open field behavior is not altered by genotype or probiotic treatment.** (A-B) (A,B) Schematic of the open field test arena shown from the side (A) and top (B). The arena was subdivided into three concentric zones representing different levels of perceived exposure: Zone 1 (center), Zone 2 (intermediate), and Zone 3 (periphery). Distances are indicated in centimeters in (A), and zones are color-coded and labeled in (B). Flies were placed in the center of the arena (Zone 1) and allowed to acclimate for 10 min prior to imaging. (C) Percentage of control and *Kdm5$^{LOF}$* flies occupying each zone following mock or probiotic treatments (all five biological replicates included). (D) Percentage of flies with gut-specific expression of *luciferase$^{RNAi}$* (control) or *Kdm5$^{RNAi}$* occupying each zone following mock or probiotic treatments (all three biological replicates included). For visualization, data are expressed as percentages; statistical analyses were performed on raw fly counts using Fisher's exact test. No significant differences in zone occupancy were detected among genotypes or treatment groups (ns).

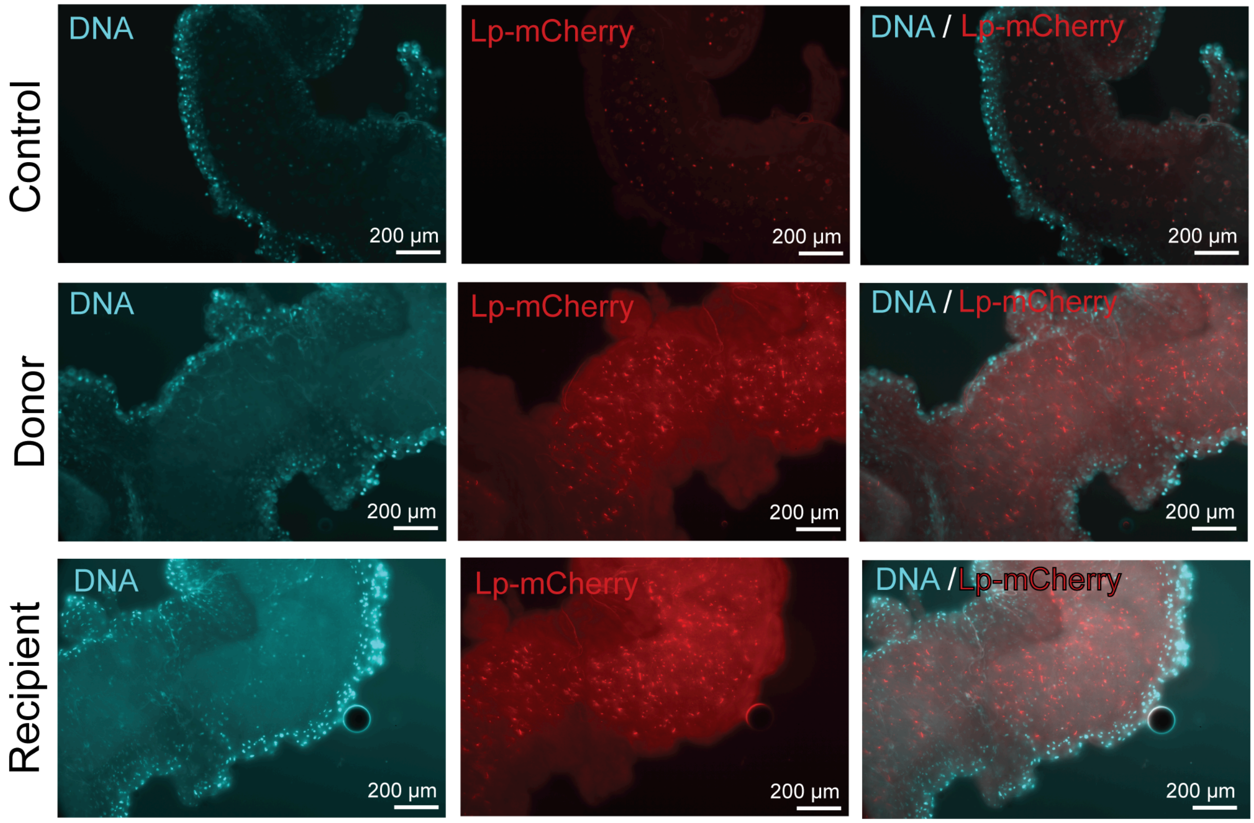


**Supplementary Fig. 3. Validation of the fecal microbiota transplantation (FMT) paradigm using fluorescently labeled *Lactiplantibacillus plantarum*.** Wild-type Canton-S female flies were either fed 5% sucrose (control) or *L. plantarum LPWF* expressing mCherry (Lp-mCherry) in 5% sucrose for 1–2 h (donor). Donor flies were subsequently transferred to fresh food without added bacteria and allowed to deposit fecal material for 3 days. Age-matched recipient flies were then placed in donor-conditioned vials and allowed to ingest fecal material for 24 h. Following treatment, flies were dissected, and guts were fixed and mounted for fluorescence imaging. DNA was labeled with DAPI (cyan), and *L. plantarum–mCherry* was detected in the red channel. Representative posterior midguts are shown for control, donor, and recipient conditions, demonstrating the presence of donor-derived bacteria in recipient fly intestines. Scale bars, 200 µm.

### Supplementary References

Rera M, Bahadorani S, Cho J, Koehler CL, Ulgherait M, Hur JH, Ansari WS, Lo T Jr, Jones DL, Walker DW. Modulation of longevity and tissue homeostasis by the Drosophila PGC-1 homolog. Cell Metab. 2011 Nov 2;14(5):623-34. doi: 10.1016/j.cmet.2011.09.013. PMID: 22055505; PMCID: PMC3238792.

Rera M, Clark RI, Walker DW. Intestinal barrier dysfunction links metabolic and inflammatory markers of aging to death in Drosophila. Proc Natl Acad Sci U S A. 2012 Dec 26;109(52):21528-33. doi: 10.1073/pnas.1215849110. Epub 2012 Dec 12. PMID: 23236133; PMCID: PMC3535647.

Martins RR, McCracken AW, Simons MJP, Henriques CM, Rera M. How to Catch a Smurf? - Ageing and Beyond… In vivo Assessment of Intestinal Permeability in Multiple Model Organisms. Bio Protoc. 2018 Feb 5;8(3):e2722. doi: 10.21769/BioProtoc.2722. PMID: 29457041; PMCID: PMC5812435.

Soibam B, Goldfeder RL, Manson-Bishop C, Gamblin R, Pletcher SD, Shah S, Gunaratne GH, Roman GW. Modeling Drosophila positional preferences in open field arenas with directional persistence and wall attraction. PLoS One. 2012;7(10):e46570. doi: 10.1371/journal.pone.0046570. Epub 2012 Oct 10. Erratum in: PLoS One. 2012;7(10). doi: 10.1371/annotation/2ccf0a7e-4f7e-47e3-9aa2-b946fbf698b7. PMID: 23071591; PMCID: PMC3468593.

Soibam B, Mann M, Liu L, Tran J, Lobaina M, Kang YY, Gunaratne GH, Pletcher S, Roman G. Open-field arena boundary is a primary object of exploration for Drosophila. Brain Behav. 2012 Mar;2(2):97-108. doi: 10.1002/brb3.36. PMID: 22574279; PMCID: PMC3345355.

Mejías K, Robles G, Martínez Z, Torres A, Algarín L, López G, Chiesa R (2016) Effects similar to anxiolysis in an organic extract of *Stypopodium zonale* on an anxiety-related behavior in *Drosophila melanogaster*. *Am J Undergrad Res* 13(4):1–8. https://doi.org/10.33697/ajur.2016.035
